# A network-based model of *Aspergillus fumigatus* elucidates regulators of development and defensive natural products of an opportunistic pathogen

**DOI:** 10.1101/2023.05.11.538573

**Authors:** Cristobal Carrera Carriel, Saptarshi Pyne, Spencer A. Halberg-Spencer, Sung Chul Park, Hye-won Seo, Aidan Schmidt, Dante G. Calise, Jean-Michel Ané, Nancy P. Keller, Sushmita Roy

## Abstract

*Aspergillus fumigatus* is a notorious pathogenic fungus responsible for various harmful, sometimes lethal, diseases known as aspergilloses. Understanding the gene regulatory networks that specify the expression programs underlying this fungus’ diverse phenotypes can shed mechanistic insight into its growth, development, and determinants of pathogenicity. We used eighteen RNA-seq datasets (seventeen publicly available and one previously unpublished) of *Aspergillus fumigatus* to construct a comprehensive gene regulatory network resource. Our resource, named GRAsp (**G**ene **R**egulation of ***Asp****ergillus fumigatus*), was able to recapitulate known regulatory pathways such as response to hypoxia, iron and zinc homeostasis, and secondary metabolite synthesis. Further, GRAsp was experimentally validated in two cases: one in which GRAsp accurately identified an uncharacterized transcription factor negatively regulating the production of the virulence factor gliotoxin and another where GRAsp revealed the bZip protein, AtfA, as required for fungal responses to microbial signals known as lipo-chitooligosaccharides. Our work showcases the strength of using network-based approaches to generate new hypotheses about regulatory relationships in *Aspergillus fumigatus*. We also unveil an online, user-friendly version of GRAsp available to the *Aspergillus* research community.

## INTRODUCTION

*Aspergillus fumigatus* is an environmentally and medically relevant filamentous fungus found worldwide. While it is considered a soil-dwelling mold, living as a saprotroph and taking part in nutrient cycling, the airborne conidia’s small size and abundance lend to its notorious reputation as a lethal opportunistic pathogen (1–3). If the fungus can bypass a person’s immune system, such as in immunocompromised individuals, it can cause various respiratory and invasive diseases. Invasive aspergillosis is the most serious of these diseases (4, 5) and has a high mortality rate (2, 6, 7). The fungus also causes a secondary infection of COVID-19, known as CAPA (COVID-19-associated pulmonary aspergillosis), that leads to an estimated mortality rate of ca. 50% (8).

The global success of *A. fumigatus* in diverse environments, from decaying organic matter to immunologically compromised patients, is reflected in significant shifts in programmed gene expression. Identification of gene regulatory networks (GRNs) that define regulatory relationships between regulatory proteins and target genes and drive these expression patterns, hold promise in elucidating key genes and pathways required for virulence or interactions in different environments. There exist at least seven studies in *Aspergillus fumigatus* that examined different sets of regulatory relationships involved in specific biological processes (9–16). For instance, Guthke et al. studied the regulatory mechanism(s) that allows *A. fumigatus* to adapt to a dramatic temperature shift during the infection process; in their study they used an inferred GRN to hypothesize that *erg11*, a gene necessary for ergosterol synthesis, modulates the expression of genes encoding heat shock proteins and the allergen RPL3 in response to a temperature shift of 30 °C to 48 °C(9). In another instance, Linde et al. inferred and experimentally validated the role of the transcriptional regulator SrbA in iron homeostasis (16). While these studies have demonstrated how GRNs can capture regulatory mechanisms, they have focused on a handful of regulators using small datasets. Uncovering genome-scale GRNs within a eukaryotic organism is a significant experimental challenge due to the sheer magnitude of experiments required to comprehensively identify and verify regulatory relationships (17, 18). Computational methods for inferring GRNs, which leverage large-scale gene expression data (i.e. mRNA transcriptomic profiles) to reverse engineer GRN structure, offer a popular and cost-effective technique to provide an initial view of the GRN that can be experimentally verified. For example, Acerbi *et al.* applied a computational method using the transcriptome of several *A. fumigatus* genes to infer the regulatory connections involved in tryptophan synthesis (11). Although these studies have shed light on GRNs in *A. fumigatus*, the datasets have been limited to specific environmental conditions (e.g., tryptophan, temperature). Our goal in this study was to construct a comprehensive genome-wide GRN of *A. fumigatus* and provide a user-friendly interface to query this GRN, enabling the generation of testable hypotheses of *A. fumigatus* molecular responses to diverse environmental conditions.

To address this goal, we applied a computational GRN inference algorithm called MERLIN-P-TFA (Siahpirani et al. in preparation, 2023, https://github.com/Roy-lab/MERLIN-P-TFA) on a curated set of publicly available and in-house RNA-seq profiles of *A. fumigatus*. This program uses a probabilistic graphical modeling approach (19) to infer a GRN (18). Moreover, MERLIN-P-TFA groups genes into modules where genes belonging to the same module have similar regulatory programs. MERLIN modules reflect small regulatory programs and provide insight into regulators driving these pathways. The MERLIN-P-TFA inferred network successfully recapitulated known regulatory relationships for multiple processes, including ergosterol biosynthesis, iron homeostasis, and secondary metabolite regulation. Further, using the network predictions of MERLIN-P-TFA, we successfully identified one transcription factor involved in mediating *A. fumigatus* response to microbial signals called lipo-chitooligosaccharides (LCOs) and one transcription factor regulating the synthesis of the bioactive toxin gliotoxin. We have created a user-friendly online resource named GRAsp (**G**ene **R**egulation of ***Asp****ergillus fumigatus*, grasp.wid.wisc.edu) that allows for visualization and exploration of *A. fumigatus’* predicted GRN.

## MATERIAL AND METHODS

### Data acquisition and preprocessing

We obtained raw fastq reads from 17 previously published and one newly generated bulk RNA-seq datasets (**Table S1**)(7, 15, 20–34). The adapter sequences were clipped with Trimmomatic (version 0.32, settings 2:30:10:2:keepBothReads LEADING:5 TRAILING:5 MINLEN:36) (35). Subsequently, the transcripts were counted with RSEM (36) using *A. fumigatus* strain Af293’s reference transcriptome ASM265v1.49 (ftp://ftp.ensemblgenomes.org/pub/fungi/release-49/gff3/aspergillus_fumigatus/Aspergillus_fumigatus. ASM265v1.49.gff3.gz). RSEM produced a transcripts per million (TPM) matrix for each dataset. In each TPM matrix, the rows represent the genes, and the columns represent the samples in that dataset. Finally, we log-transformed and quantile-normalized each TPM matrix. These initial matrices were given as input to the batch effect correction step.

### Batch correction

Each TPM matrix was zero-mean transformed, i.e., the mean of each gene’s expression was subtracted from its expression value in each sample. As a result, each gene had a mean expression of zero in every TPM matrix after the transformation. Subsequently, we performed principal component analysis (PCA) on the transformed matrices. We generated PCA plots using the combined matrix before and after zero-mean transformation (**Figure S5, S6, and S7**). PCA components after zero-mean transformation demonstrated that the variation between datasets was less prevalent than variation between experimental conditions. To further verify that batch correction was sufficient to remove variation between datasets we generated correlation and gene expression heatmaps. We identified one dataset, namely PRJEB2987, which was observed to have outlier samples. We observed seven of the samples, S1-S4 and samples from the third replicate (rep3), with unexpected correlation structure and expression (**Figure S8 and S9)**. We removed these seven samples from our dataset. The final expression matrix contained the values of 9,859 genes across 294 samples.

### Inference of the gene regulatory network

To infer a GRN from the expression matrix, we applied a network inference method named MERLIN-P-TFA (Siahpirani et al., 2023, https://github.com/Roy-lab/MERLIN-P-TFA). This method extends our previous method, MERLIN-P, by estimating hidden TF activity levels from a noisy input prior network (37). This algorithm takes three inputs: (a) an expression matrix, (b) a list of regulators, and (c) a prior network. Only the genes mentioned on the input list of regulators are allowed to have outgoing edges in the inferred network.

#### Preparation of the list of regulators

We curated five lists of regulators and combined them to produce the final list of regulators. The lists are: (a) the predicted TFs for A. *fumigatus* strain Af293 in the JGI MycoCosm database, (b) the predicted TFs in Furukawa et al. (2020), (c) the signaling genes that are annotated for GO terms “G-protein coupled receptor protein signaling pathway” (GO:0007186) and “Calcium-mediated signaling” (GO:0019722) in the JGI MycoCosm database, (d) the list of G-protein coupled receptors (GPCRs) found in (38–40), and (e) the genes annotated for “Signal transduction” (GO:0007165) in the AspGD gene ontology database (38–41). The combined list consists of 820 distinct regulators (**Figure S1**(40)). In Furukawa et al., the TF names correspond to A. *fumigatus* strain A1160; hence, we used OrthoFinder to find the orthologous gene names for strain Af293 (39, 42, 43). See Supplementary File **S1_regulators.xlsx** for a detailed description of each sublist.

#### Construction of the prior network

MERLIN-P-TFA utilizes a prior network to guide its network inference task by incorporating prior knowledge on an edge when available. We utilized putative TF binding site sequence motifs to construct the prior network. We downloaded 628 (DNA-sequence) motifs corresponding to 99 TFs in *A. fumigatus* from the CIS-BP database (44). Additionally, we used four previously characterized *A. fumigatus* motifs corresponding to two TFs (45, 46); for brief descriptions, see **Supplementary Method S1**(45, 46). We scanned the A. *fumigatus* reference genome (ASM265v1 release 49) for each motif to determine the genome-wide coordinates of the motif instances on the genome. Each set of coordinates was then mapped to a gene if the coordinates overlapped a region 10 Kbp upstream and 1 Kbp downstream of the gene transcription start site (TSS). Suppose motif M has two sets of coordinates that were mapped to genes A and B, respectively, and TF X can bind to motif M. In that case, two edges were included in the prior network, one from X to A, another from X to B (**Figure S2**)(47). Moreover, we assigned each edge a PIQ score representing the likelihood of the motif appearing on the binding site of the target gene (47). The scores were calculated with the R package “Biostrings” (48, 49). We sorted the candidate binding sites by their scores for each motif and retained only the top 100,000 sites. For motifs with less than 100,000 candidate binding sites, we retained all the sites. The resultant prior network was extremely dense (containing 936,557 edges). Hence, we restricted its density to be 20% of the total number of possible edges (0.2 x Number of TFs in the prior network, i.e., 96 × Number of target genes, i.e., 9,859), which corresponded to 189,292 edges. Finally, we converted the range of the edge scores to be between 0 and 1 by applying percentile ranking to them. The highest score was 1, and the lowest score was 0.000101688 (Supplementary File **S2_prior_network.xlsx**).

#### Network inference with MERLIN-P-TFA

MERLIN-P-TFA performs the network inference in two steps: First, it estimates the transcription factor-level activities (TFA) of the regulators present in the prior network. Second, it infers the edges between the regulators and the target genes by regressing the estimated TFA profiles of the regulators present in the prior network as well as the gene expression profiles of all the regulators to the gene expression profiles of the candidate target genes (**Figure 1**).

**Figure 1.**
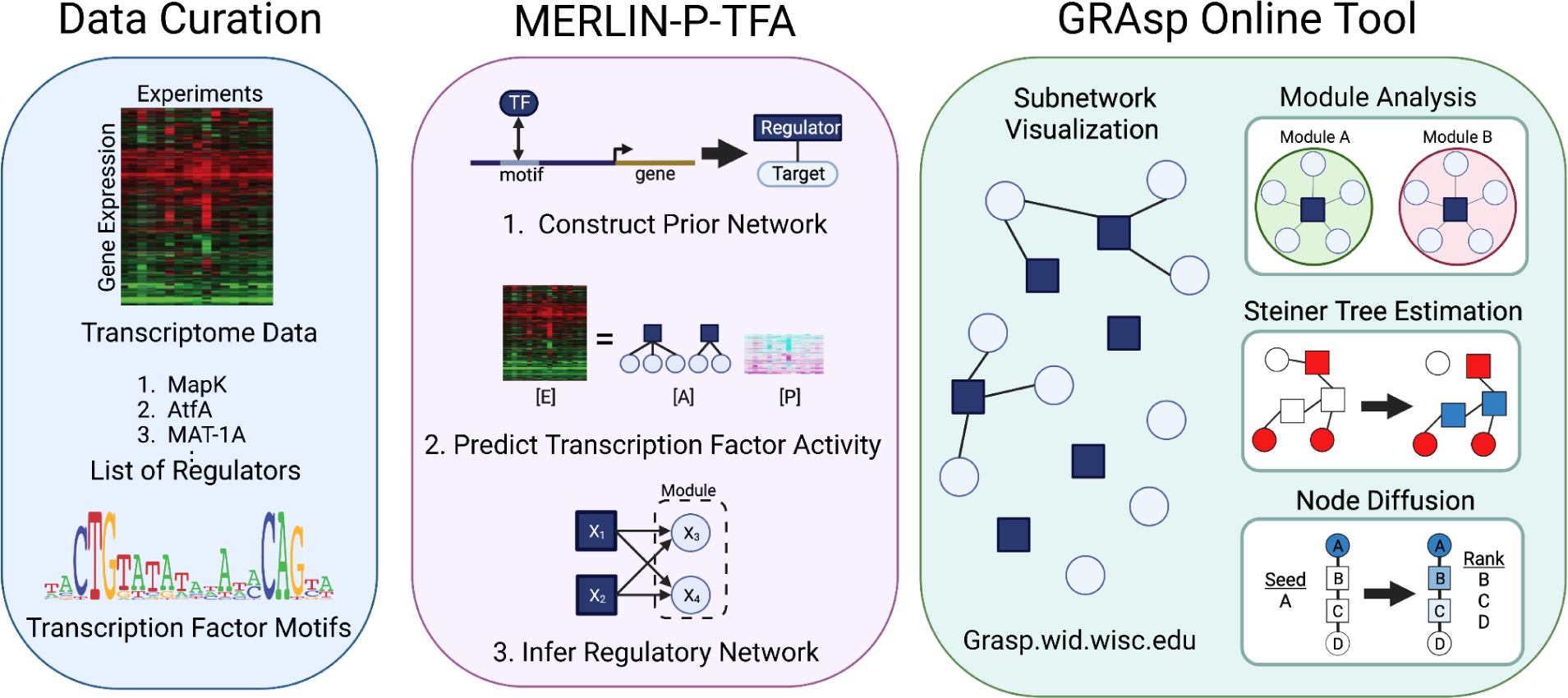
The Network-based Modeling and Analysis Workflow. ***Data Curation:*** We curated 18 *A. fumigatus* bulk RNA-seq profiles and combined them to produce a batch-corrected transcriptome data matrix of 9,859 genes across 294 experiments. We also prepared a list of 820 putative regulator genes and 632 TF binding site sequence motifs. ***MERLIN-P-TFA:*** We used our MERLIN-P-TFA method to infer a gene regulatory network from the curated expression data. MERLIN-P-TFA takes as additional input a prior network which was constructed based on the TF motif information. Given the prior network, the list of regulators, and the transcriptome data matrix, MERLIN-P-TFA first estimated a transcription factor activity (TFA) matrix. This matrix represents the estimated TF-level activity of the regulators (that are present in the prior network) across all the experiments. Subsequently, MERLIN-P-TFA combined the TFA matrix with the transcriptome data matrix to infer the gene regulatory network. ***GRAsp Online Tool:*** GRAsp is an online visualization framework for interacting with the inferred regulatory network. GRAsp offers various network analysis techniques, such as gene module analysis, Steiner tree estimation, and node diffusion, to recapitulate known pathways and hypothesize novel gene regulatory activities (some of which were further experimentally validated). Created with Biorender.com.

In the first step, MERLIN-P-TFA estimates the TFA matrix. The TFA matrix is a regulators-by-samples matrix where the (i, j)^th^ entry denotes the estimated TF-level activity (50–52) of the regulators (that are present in the prior network) across all the samples (in the expression matrix). To estimate the TFA matrix, *P*, MERLIN-P-TFA solves the minimization problem 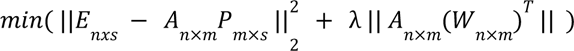 where *E* is the given expression matrix, *n* is the number of target genes, *s* is the number of samples, *A* (called the “connectivity matrix” which is derived from the given “prior network”) is the prior knowledge on the regulatory edges between the regulators and target genes, *m* is the number of regulators, and *W* is the prior edge-penalty matrix; *W* (i, j) represents the prior penalty on the edge from the j^th^ regulator to the i^th^ target gene calculated as (1 - the confidence of the edge in the prior network). The regularization term 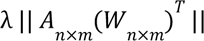 is important for handling noise in the connectivity matrix (Siahpirani et al. in preparation, 2023, https://github.com/Roy-lab/MERLIN-P-TFA). The level of regularization depends upon the value of λ, the higher the level of regularization. In this work, we used λ = 0. 1. λ; the higher Once the TFA is estimated, MERLIN-P-TFA utilizes the original MERLIN-P method to infer a GRN. Briefly, MERLIN-P represents the relationships between regulators and target genes as a probabilistic graphical model (PGM) known as a “dependency network.” Given a dataset *D*, MERLIN-P infers a network *G* by solving a set of regression problems. Additionally, MERLIN-P groups genes into modules where the genes having similar regulatory programs are assigned to the same module.

For each gene, MERLIN-P-TFA infers two items: (a) a set of directed edges from its predicted regulators to the gene itself and (b) a module assignment. We ran the TFA estimation step 50 times by randomly initiating the connectivity matrix “A” every time. This resulted in 50 estimated TFA matrices. We combined these matrices into a single TFA matrix by taking their mean. Subsequently, we produced a large matrix by appending the estimated TFA matrix to the gene expression matrix. This large matrix contained the expressions of 9,859 genes and the estimated TFAs of 30 regulators across all 294 samples. Then we generated 100 random subsamples of the large matrix through random sampling without replacement. Each subsample matrix contained 147 samples, i.e., half the number of samples in the large matrix. We ran MERLIN-P-TFA separately on each of these 100 subsample matrices. This results in 100 distinct GRNs, each having a unique module assignment (i.e., gene to module map). We combined these 100 GRNs into a single GRN by taking the union of their edges with each edge assigned a confidence score that reflects how many times the edge had appeared in the previous 100 GRNs. For downstream analyses, we only considered the edges having a confidence score of 80% or above. We also inferred a consensus module assignment requiring genes to be in the same module at least 30 out of 100 module assignments.

### Interpretation of MERLIN-P-TFA network and modules

To interpret the results of MERLIN-P-TFA and the generation of new hypotheses, we developed three strategies based on module enrichment and graph-theoretic analyses. All three methods accept input gene lists representing known gene sets, such as from a Gene Ontology (GO) database or gene list from an independent differential expression analysis experiment.

#### Interpretation of MERLIN modules based on enrichment analyses

MERLIN-P-TFA infers a per-gene module-constrained network. MERLIN-P-TFA modules are sets of genes with similar expression profiles and have similar, although not identical, regulatory programs. To interpret the MERLIN-P-TFA modules, we tested them for enrichment of GO terms using a hypergeometric test with FDR correction for multiple hypotheses (53). A module was labeled with a GO term if a significant number of genes in this module were enriched for that GO term (FDR<0.05). In addition to the GO enrichment analysis, we tested the modules for enrichment of gene sets identified from targeted experiments in *Aspergillus*, e.g., differentially expressed genes from a gene perturbation experiment. A similar FDR-corrected p-value threshold was used to test for enrichment of such gene sets in the modules. Modules were also useful for determining important regulators of specific gene programs. Since MERLIN-P-TFA does not require all genes in the same module to have the same set of regulators, we further use enrichment tests to determine the important regulators of a module using a similar hypergeometric test with FDR correction (FDR<0.05).

#### Approximate Steiner Tree Construction for Identification of Related Regulatory Mechanisms

While modules capture part of a regulatory program indicated by the gene expression pattern, an entire regulatory program likely spans multiple modules as well as individual genes that may not be co-expressed with many other genes but are still part of the GRN. Thus we developed a second method to identify important pathways connecting a list of genes within a program to a set of modules and additional genes. To accomplish this task, we implemented a Steiner tree-based method to find approximately minimal trees connecting any user-provided set of genes. Briefly, the Steiner tree problem on unweighted, undirected graphs is defined as given a graph *G* = (*V, E*), where *V* denotes the set of vertices and *E* denotes the set of edges and a set of terminals *T*, finds a subgraph G* = (*V*\*, *E*\*) that spans *T* such that the cardinality of *E*\* is minimized. Unfortunately, the Steiner tree problem for more than two terminals is computationally intractable; thus, an approximation of such a tree is generated (54). The closest two genes, based on shortest path, from the list are connected to generate the approximation and form an undirected subgraph. The next closest gene is then iteratively added to the new subgraph until all terminal genes in T are added. In our application of the Steiner tree approach, the terminal nodes come from an input gene list generated from differential expression or similar analyses. To guarantee algorithm convergence, genes from the gene list are only considered if they are contained within the largest connected component of the predicted regulatory network. Although the approximate Steiner tree is not unique, the tree can identify hub genes that are important for the regulation of multiple different components of the gene regulatory programs.

#### Ranking of Gene Importance via Laplacian Kernel Node Diffusion

Our third method of determining the relationship between a specific gene pathway and the predicted regulatory network is based on a gene prioritization scheme based on diffusion on a graph (55). This strategy has been used to predict new genes for pathways as well as to examine the impact of genetic mutations in one gene on other genes connected via a protein-protein interaction network (55). In this approach, a graph kernel is used to define global similarity between nodes in the input set to all other nodes in the graph. In our case, we used the Laplacian graph kernel defined as 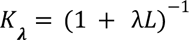, where λ is a hyper-parameter specifying the bandwidth of the kernel function and *L* is the graph Laplacian matrix defined as *L* = *D* − *A*, where *D* is the diagonal matrix representation of each node degree and *A* is the symmetric graph adjacency matrix. Diffused scores for all genes are defined by *V* = *K*_λ_ *S*, where *S* are the initial scores specified by the user. In our setup, the input gene set corresponds to genes from a pathway of interest or differentially expressed genes, and the graph is the MERLIN-P-TFA inferred GRN. The input gene set is initialized with uniform scores of 1 or based on additional data, such as the change in expression in two conditions of interest. Diffused scores are then used to prioritize genes based on their connectivity to the input gene set which can indicate particular gene pathways of interest.

### Fungal Strains and culture conditions

*Aspergillus fumigatus* strains used in this study are listed in **Table S2**(56–60). All strains were grown at 25 °C for 9 days in the dark on solid glucose minimal medium in sterile 100 x 15 mm Petri dishes (61). Ten thousand spores were used for the initial inoculum. Conidia were collected in sterile 0.01% Tween 80, filtered through a 40 μm nylon cell strainer, then quantified using a hemocytometer. For long-term storage, conidia were maintained as glycerol stocks at −80 °C.

### Generating rogA mutant strains

To create the *rogA* (AFUA_3G11990) deletion strain (TSCP2), its 1.4 kb 5’ flanking region and 1.0 kb 3’ flanking region were amplified by PCR from the genomic DNA from *A. fumigatus* strain AF293. The *A. parasiticus pyrG* gene was used as a selective marker and was amplified by PCR from the pJW24 plasmid (62). To make the *rogA* overexpression strain (THWS25), 1.0 kb of the gene’s 5’ flanking region and the entire 1 kb *rogA* region, starting with ATG, were amplified from Af293 genomic DNA. The *pyrG* linked with the *A. nidulans gpdA* promoter was amplified from the pJMP9 plasmid (63).

These fragments were fused by double-joint PCR, respectively (64). About 25 μL of Sephadex® G-50 purified third-round PCR product was used to transform TFYL80.1 for both the deletion and overexpression mutants. All of the fungal transformations were done using the polyethylene glycol (PEG)-based method previously described (64). Both the deletion and overexpression mutants were confirmed by PCR and Southern blot (**Figure S3**).

### Metabolite Extraction

GMM cultures of WT, Δ*rogA*, and OE::*rogA* strains were grown (culture conditions described above) and lyophilized for 3 days, in quadruplicate. The dehydrated agar and biomass were then crushed to powder. Each sample was extracted with 25 mL of 100% methanol and passed through filter paper. Extracts were reduced by air drying, weighed, and resuspended in 100% methanol at a final concentration of 1 mg/mL. Two authors performed these experiments independently.

### UHPLC–HRMS and UHPLC–MS/MS analyses

UHPLC–HRMS was performed on a Thermo Scientific Vanquish UHPLC system connected to a Thermo Scientific Q Exactive Hybrid Quadrupole-Orbitrap mass spectrometer operated in positive ionization mode. A Waters Acquity UPLC BEH-C18 column (2.1 × 100 mm, 1.7 μm) was used with acetonitrile (0.1% formic acid) and water (0.1% formic acid) at a flow rate of 0.2 mL/min. A screening gradient method was implemented as follows: Starting at 10% organic for 5 min, followed by a linear increase to 90% organic over 20 min, another linear increase to 98% organic for 2 min, holding at 98% organic for 5 min, decreasing back to 10% organic for 3 min, and holding at 10% organic for the final 2 min, for a total of 37 min. Ten μL of each sample was injected into the system for the analysis. Gliotoxin was identified by comparison with a standard purchased from Cayman Chemical (Ann Arbor, MI, USA). An analog of gliotoxin, BMgliotoxin, was predicted by the analysis of UHPLC–MS/MS data through SIRIUS ver.5.5.7. The relative quantification of two compounds was calculated based on intensities obtained from UHPLC–MS/MS.

### Analysis of secondary branching and germination in *A. fumigatus*

To look at the number of secondary branches in *A. fumigatus*, 1×10^6^ conidia per mL of *A. fumigatus* strains was inoculated in GMM broth treated with either 10^-8^ M sulfated C16:0 LCO (CERMAV) or the negative control, 0.005% ethanol (61). 100 μL of inoculated and treated broth was dispensed into a well in a 96-well flat bottom plate in triplicate. The plate was incubated at 37 °C for 11 hours, followed by imaging every 15 minutes over 3 hours using a Nikon TI inverted microscope with a 40X objective. Each frame captures the growth of one hypha, and ten frames were taken per well. The number of secondary branches per apical hyphae was counted using NIS-Elements AR Analysis Version 4.30.

To quantify the germination of conidia, 1×10^5^ spores per mL of *A. fumigatus* strains were inoculated in GMM broth treated with either 10^-8^ M sulfated LCOs (LCO-IV (C:16,S)) or 0.005% ethanol (negative control). 1 mL of inoculated and treated broth was dispensed into a well in 24-well flat-bottom plates and incubated at 37°C for 3 hours. After 3 hours, 10 different areas in the well were captured as frames using a 40X objective. Each frame contained 20-30 conidia and was photographed every hour for 12 hours. Germination of conidia was counted using NIS-Elements AR Analysis Version 4.30.

### RNAseq extraction from *A. fumigatus zfpA* mutants

For mycelial RNA, *A. fumigatus* wild type, Δ*zfpA*, and OE::*zfpA* conidia were inoculated at 10^6^ spores per mL in liquid glucose minimal media and incubated at 37°C shaking at 250 RPM overnight (60). For each sample, the total tissue was combined from two 50mL cultures flash frozen and lyophilized. Total RNA was extracted using QIAzol Lysis Reagent (Qiagen) according to the manufacturer’s instructions with an additional phenol:chloroform:isoamyl alcohol (24:1:1) extraction step before RNA precipitation. For conidial RNA, 10^6^ WT, Δ*zfpA*, or OE::*zfpA* conidia were inoculated into 8mL of molten GMM top agar (0.75% agar) and overlaid onto cooled GMM plates containing 15mL bottom media (1.5% agar). Plates were incubated at 37°C for 3 days before harvesting in 0.01% Tween 80 solution. Spore suspensions were filtered twice through two layers of sterile miracloth and once through 40µm cell strainers (Falcon) to remove hyphal fragments. Filtered conidia were pelleted at 1000 x g for 10 minutes and resuspended in 200µL 0.01% Tween 80. Concentrated conidia were lysed by bead beating for 5 minutes with 0.5mm zirconia/silica beads after the addition of QIAzol Lysis Reagent and chloroform. Total RNA extraction was completed per manufacturer’s instructions with an additional phenol:chloroform:isoamyl alcohol (24:1:1) extraction step before RNA precipitation. Total RNA from both mycelial and conidial samples were further cleaned up using RNeasy Mini Kit with on-column DNase digestion (Qiagen) per manufacturer’s protocol. RNA integrity was tested via nanodrop, gel electrophoresis, and the Agilent 2100 Bioanalyzer. Library preparation and RNA sequencing were performed by Novogene, Inc. using the TruSeq Stranded mRNA Library Prep Kit and Illumina Novaseq 6000 Platform.

### Statistical Analysis

#### Gliotoxin Mass-spectrometry

Relative abundance of BMgliotoxin and gliotoxin were measured by two authors in independent experiments. Normalization was performed for each author independently by dividing all relative abundance measurements with respect to the mean of the WT condition. After normalization, data was pooled between authors (total *n*=8 per strain). Statistical analyses between wild type and the genetically modified strains was performed using an Unpaired t-test (Graphpad Prism v9.4.1).

#### Network Diffusion utilizing differentially expressed genes of LCO treated samples

For diffusion analysis, we used a list of differentially expressed genes (DEGs), published in Rush et al. 2020, of *A. fumigatus* 30 minutes after treatment with LCOs (**Table S1**, Serial number 16)(33). Diffusion was performed using Laplacian kernel diffusion with hyperparameter, λ=10 and the absolute log fold change of the DEGs. Regulators with at least 5 targets were sorted to determine importance with respect to the LCO treated condition.

#### AtfA mutant phenotype quantification in LCO treated samples

Secondary branching and percent germination were quantified as described above. Germination studies had a *n=*16 total for WT; △*atfA, n*=19, OE::*atfA, n*=17. Branching studies had a *n=*60 total for WT; △*atfA, n*=59, OE::*atfA, n*=57. Statistical analyses comparing the phenotypic traits in *AtfA* relative wildtype after treatment with LCO were performed using an Unpaired t-test utilizing Graphpad Prism (v9.4.1).

## RESULTS

### Genome-scale Gene Regulatory Network of *A. fumigatus* identified by MERLIN-P-TFA

MERLIN-P-TFA generated a genome-wide gene regulatory network (GRN) based on eighteen RNA-seq datasets of multiple *A. fumigatus* strains in a variety of conditions **(Table S1, Figure 1**). Our network, which consists of 80% confident edges, contains 7,422 regulatory edges. Each regulatory edge connects one of 669 regulators to one of the 5274 target genes. Our regulator set includes 12 signaling proteins (**Figure S1**) and 30 regulators with motifs. MERLIN-P-TFA estimated the TFA of these 30 regulators. The gene encoding the mating type protein MAT1-2 and the stress response protein AtfA (incorporated based on TFA) are the top two regulators of the network, regulating 216 and 155 target genes, respectively (**Figure 2B**). Furthermore, MERLIN-P-TFA identified 164 modules with at least five genes. Jointly they contain 3,381 genes. To interpret the biological relevance of these modules, we performed a GO enrichment analysis. Among the 164 modules, 74 modules were enriched with at least one GO term (Supplementary file **S3_module_details.xlsx**).

**Figure 2.**
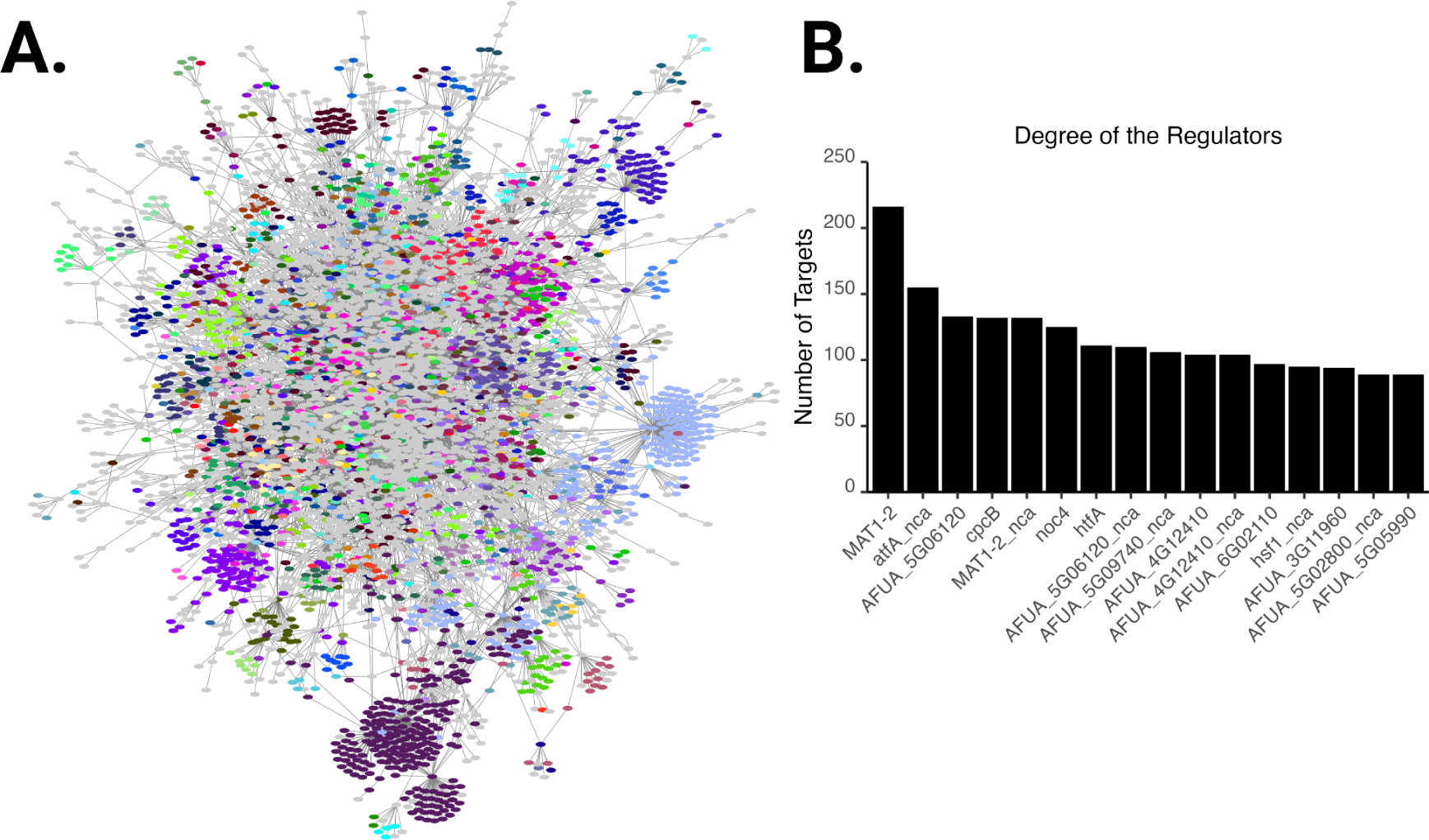
*A. fumigatus* gene regulatory network predicted by MERLIN-P-TFA. A) Force-directed layout of the inferred network where genes belonging to the same module are depicted with the same color. Visualization of the network was performed using Cytoscape (v.3.8.2) (65). B) Predicted major regulators ordered by their number of targets. The regulators having the “_nca” suffix indicates that the estimated TFA of the regulator was included in the network based on its TFA. For example, “atfA_nca” represents that “atfA” was selected as a regulator based on its TFA. The “_nca” suffix refers to the regularized “network component analysis” technique used by MERLIN-P-TFA for estimating the TFA profiles of the regulators (Siahpirani et al., 2023, https://github.com/Roy-lab/MERLIN-P-TFA). Figure prepared with Biorender.com

### GRAsp: a visualization framework for interpretation and hypotheses generation of the ***Aspergillus* GRN**

Genome-wide regulatory networks are difficult to interpret due to the large number of predicted interactions between regulators and target genes. To aid in interpreting and analyzing our MERLIN-P-TFA inferred network, we have developed a network visualization framework, called GRAsp (**G**ene **R**egulation of ***Asp****ergillus fumigatus*), that incorporates different graph theoretic tools to examine sets of genes related to a specific process of interest. GRAsp allows users to input a list of genes or gene ontology terms, then actively generates interactive network diagrams that can be used to determine regulators of interest. The networks visualized in the GRAsp display window can be extended to include all genes in associated MERLIN modules or in the neighborhood of the list of selected genes, providing additional information about regulatory mechanisms of interest. GRAsp also incorporates a Steiner tree estimation method to connect sets across MERLIN modules. Given a list of genes of interest, the Steiner tree method connects all genes with an approximate spanning tree (**Methods**). The final approach in GRAsp allows the user to incorporate additional data into the network using node diffusion (55). For example, this feature allows users to use differential expression p-values or fold change information to prioritize regulators. The nodal gene values are assigned to each node and diffused via a Laplacian kernel (55). GRAsp then displays the top regulators related to the pathway. The network visualization panel can be customized and saved as publication-quality figures. The GRAsp network tool is publicly accessible at <u>grasp.wid.wisc.edu</u>.

### Diverse biological processes of *Aspergillus* are captured in GRAsp

We first evaluated GRAsp for its ability to recapitulate known regulatory pathways in *A. fumigatus*. Specifically, we looked at four previously characterized transcription factors and their roles in regulating different pathways: SrbA and its role in regulating hypoxia and various stress responses, ZafA and its response to zinc starvation, HapX and its response to iron starvation, and FapR’s regulation of secondary metabolites.

#### SrbA and hypoxia

The ability of *Aspergillus fumigatus* to respond to hypoxia, low iron availability, and antifungals contributes to the fungus’ pathogenicity. The response to these conditions is partly mediated by the sterol regulatory element binding transcription factors SrbA and SrbB (16, 66). Chung et al., 2014 employed ChIP-seq technology to uncover the downstream targets of these two transcription factors. Under hypoxic conditions, SrbA and SrbB regulate each other while also regulating target genes involved in adaptation to hypoxia, such as ergosterol biosynthesis (*erg1, erg11A, erg25A, erg3B, erg5*), nitrate assimilation (*niaD*, *niiA*), nitric oxygen (NO)-detoxifying flavohemoprotein gene (*fhpA*), heme biosynthesis (*hem13*), and ethanol fermentation (*alcC*) (67). The module containing *srbA*, module number 5395, recapitulated many of the known regulatory relationships (**Figure 3A, B)**. Additionally, many genes, such as *erG3A* or *niaD*, which are not predicted to be directly regulated by either *srbA* or *srbB*, are still present in the MERLIN module. Only the regulatory relationship between *srbB* and *alcA* was missed entirely. Most importantly, GRAsp inferred new genes with additional functions in these processes, such as sterol biosynthesis (*cyp51A*) and heme biosynthesis (*hem14*), as well as genes of currently unknown function (in black in **Figure 3B**).

**Figure 3.**
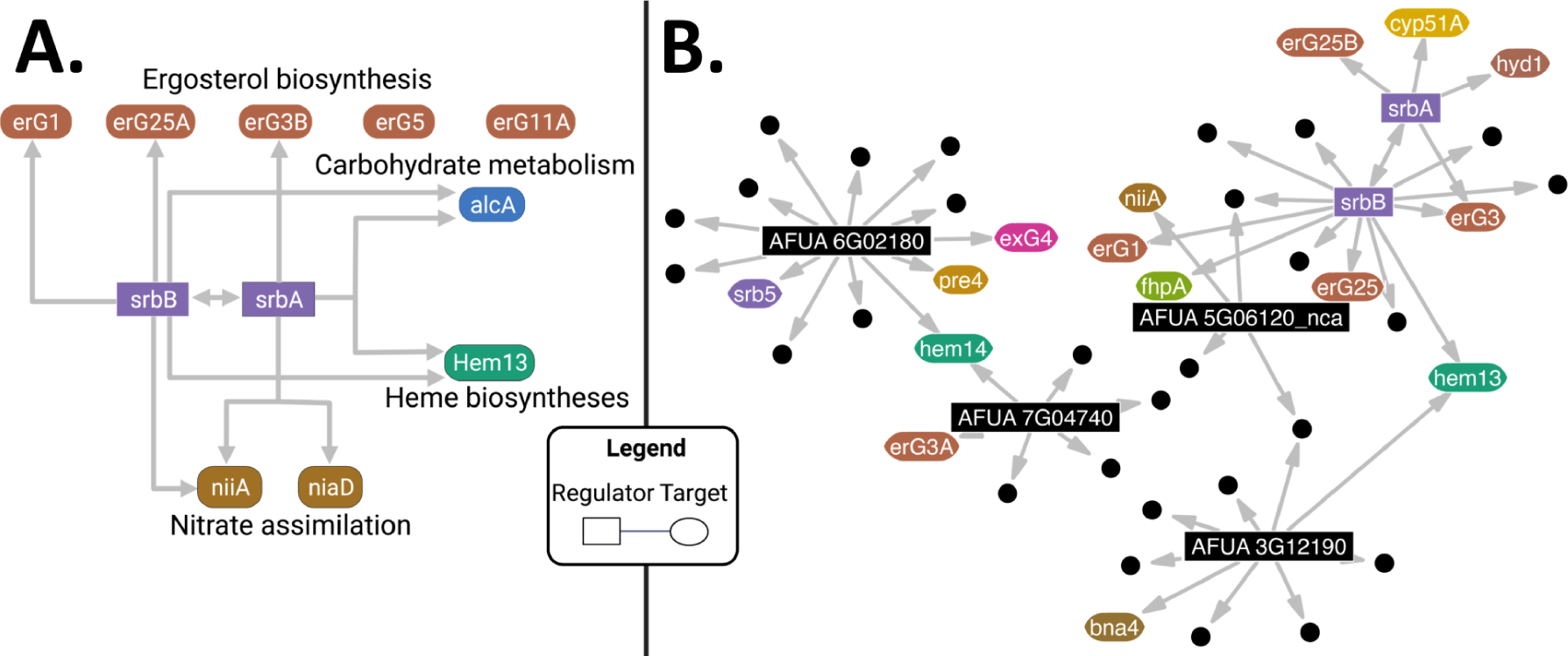
Recapitulation of SrbA targets. A) SrbA pathway inferred through ChIP-seq, reproduced from Chung *et al.* 2014 (Published under a creative commons attribution license). Genes in rectangles represent regulators; gene targets are contained in ovals. Gene colors represent each pathway. Created with Biorender.com. B) Module 5395, which includes the transcription factors *srbA* and *srbB* visualized with GRAsp. Black circular nodes represent gene targets without a gene symbol in the fungiDB database. Genes within the module that are not part of the known *srbA* and *srbB* regulatory system and are not regulated by any of the module regulators have been removed to simplify the figure. Figure prepared with Biorender.com

#### ZafA and zinc homeostasis

Zinc is important for proper growth and development in *A. fumigatus* (*68*). In circumstances where the fungus encounters low zinc environments, such as within host tissue, the acquisition of zinc ions is essential. To maintain zinc homeostasis, the transcription factor ZafA regulates *zrfA*, *zrfB*, and *zrfC* expression, which encode transporters involved in zinc uptake (68, 69). Module 5027, which is regulated by ZafA, contains *zrfA.* Additionally, *zrfB* and *zrfC* which are outside of this module are still predicted targets of *ZafA* (**Figure 4A**). The resulting subnetwork displays ZafA as the primary regulator of 15 target genes, including *zrfA*, *zrfB*, and *zrfC*. The subnetwork also reveals uncharacterized targets of ZafA, that perhaps could also be involved in zinc homeostasis. For example, AFUA_5G02010 has orthologs related to metalloendopeptidase activity and AFUA_1G14700 is related to transmembrane transport.

**Figure 4.**
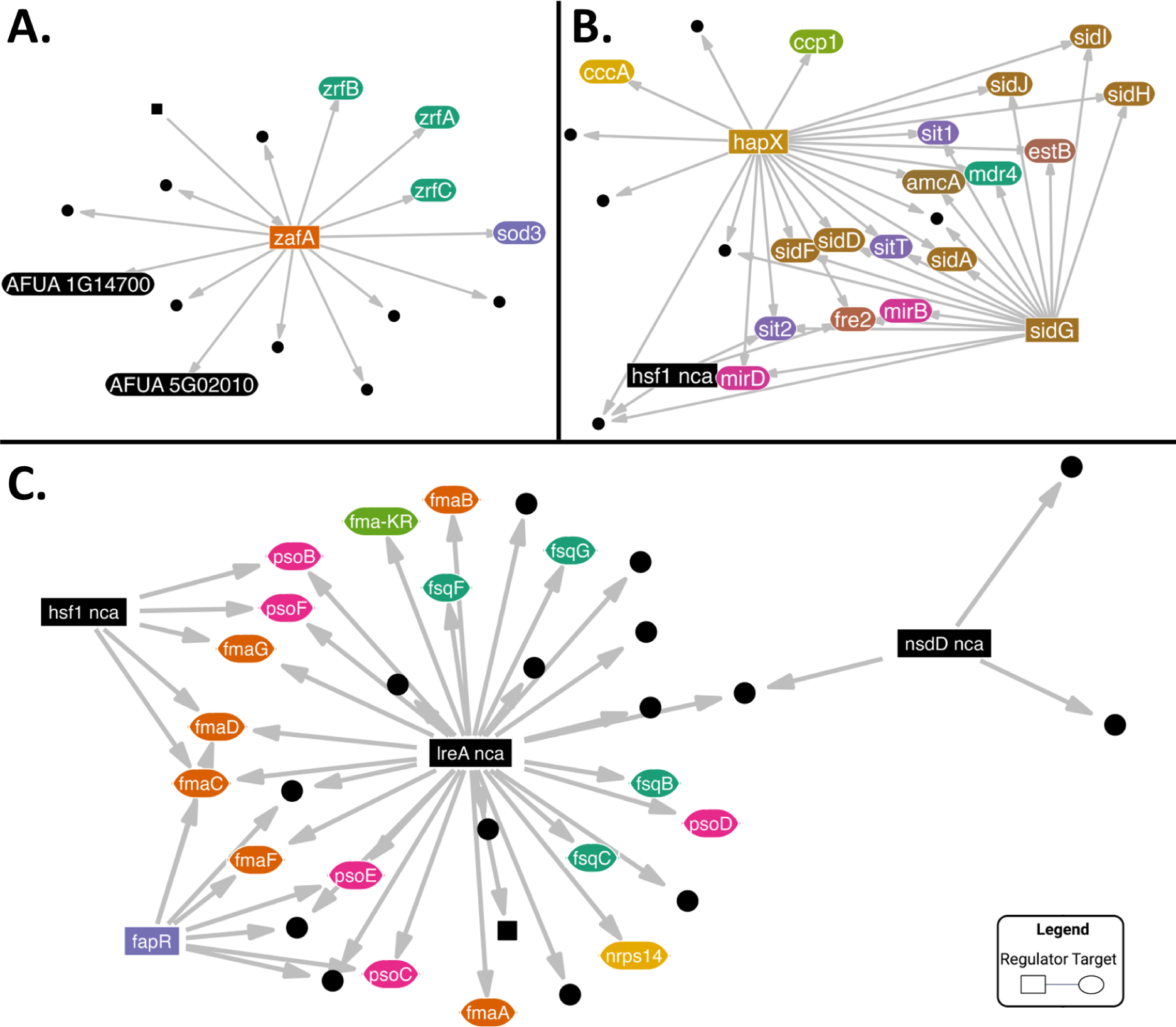
Recapitulation of known genetic pathways in GRAsp. Network representations of known genetic pathways are visualized with GRAsp. In all figures, node colors correspond to similar common name prefixes, i.e. fma, fsq, zrf, and sid. Black nodes represent gene targets without a gene symbol in the fungiDB database. Regulators are represented as rectangles and squares, targets are represented as circles and ovals. A) The regulatory targets of *ZafA* generated by selecting its neighbors. B) The modules 4901 and 5283 which contain the *hapX* gene. C) The module 5396 which contains the *fapR* gene. Orange target genes represent fumagilin biosynthesis genes (fma prefix), pink targets represent pseurotin genes (pso prefix), and green targets represent fumisoquin genes (fsq prefix). Figure prepared with Biorender.com.

#### HapX and iron starvation

Similar to zinc, iron is a critical cofactor metal in *A. fumigatus*, and processes to acquire iron under iron limiting conditions are regulated in part by the transcription factor HapX (70). Specifically, HapX plays an essential role in activating genes involved in the synthesis of specialized iron-acquiring metabolites called siderophores (*sid* genes and *estB* encoding a siderophore triacetylfusarinine C esterase), putative siderophore transporters (*sit* genes and *mirB* and *mirD*), and the metalloreductase important during iron starvation (*fre2* gene). The inferred MERLIN-P-TFA regulatory network contained modules 4901 and 5283 which are regulated by HapX (**Figure 4B**). These modules contain all of the known targets of HapX.

#### FapR and secondary metabolites

*A. fumigatus* is renowned for its arsenal of secondary metabolites (SMs) that play critical roles in various ecological settings, from pathogenesis to encounters with other microbes (71). The genes required to synthesize a SM are usually grouped together on the genome in what is known as a biosynthetic gene cluster, BGC (72). A prime example is the metabolite fumagillin. This compound, a virulence factor in invasive aspergillosis, targets methionine aminopeptidase, which removes the amino-terminal methionine residue from newly synthesized proteins (73). This functionality has led to various fumagillin applications, such as microsporicidal activity in treating honey bee hive infections (74). Interestingly *A. fumigatus* is resistant to fumagillin, presumably due to extra copies of methionine aminopeptidase genes in its genome (75).

The fumagillin BGC is unique because its biosynthetic genes are intertwined with genes of another SM, pseurotin, known for its antibacterial properties (75, 76).The gene encoding the transcription factor FapR is embedded in the intertwined cluster and FapR regulates the biosynthesis of both pseurotin *(psoF, psoG, psoB, psoA, psoC, psoD*, and *psoE)* and fumagillin (*fmaB, fmaC, fmaD, fmaG*, and *fmaA*). Querying GRAsp with *fapR* as input gene found a MERLIN-P-TFA module containing many *fma* and *pso* biosynthetic genes (**Figure 4C**). Interestingly MERLIN-P-TFA suggested a relationship between fumagillin/pseurotin production and another secondary metabolite, fumisoquin (*fsq* genes) (77). At present, the ecological role of fumisoquin is unknown.

### MERLIN-P-TFA successfully predicts a novel regulator of gliotoxin

Considering the accurate recapitulation of known regulatory pathways by MERLIN-P-TFA, we next asked if we could successfully identify new regulatory connections using our resource. We were interested in querying the regulation of the SM gliotoxin, a virulence factor in murine models of invasive aspergillosis and a potent antifungal (78, 79). Previous work has shown that the gliotoxin BGC (*gli* BGC) is regulated by an in-cluster transcription factor GliZ (**Figure 5A**); however, how *gliZ* is regulated is largely unknown (80). Using *gliZ* as the query gene, GRAsp revealed a module enriched in most of the *gli* genes (**Figure 5B**). The module was predicted to be regulated by an uncharacterized gene, *AFUA_3G11990*, encoding for a GAL4 type C6 transcription factor. We deleted and overexpressed *AFUA_3G11990* to see how it affected the production of gliotoxin and its derivative bis(methylthio)gliotoxin (**Figure 5C**), both of which can be detected in human serum and are potential diagnostic indicators of *Aspergillus* infections (81). In the two *AFUA_3G11990* deletion mutant siblings, Δ*rogA.1 and* Δ*rogA.2*, we observed a significant increase in gliotoxin and no change to bis(methylthio)gliotoxin. Furthermore, we observed a significant decrease in both gliotoxin production and bis(methylthio)gliotoxin in the two *AFUA_3G11990* overexpression mutants siblings,OE::*rogA.1 and* OE::*rogA.2*,suggesting that AFUA_3G11990 is a negative regulator of the *gli* BGC, possibly through regulation of *gliZ* expression. Thus, we named AFUA_3G11990 as *rogA* for **r**egulator **o**f **g**liotoxin. The repressive regulatory relationship between AFUA_3G11990 was also observed in the normalized gene expression matrix (**Figure S4**).

**Figure 5.**
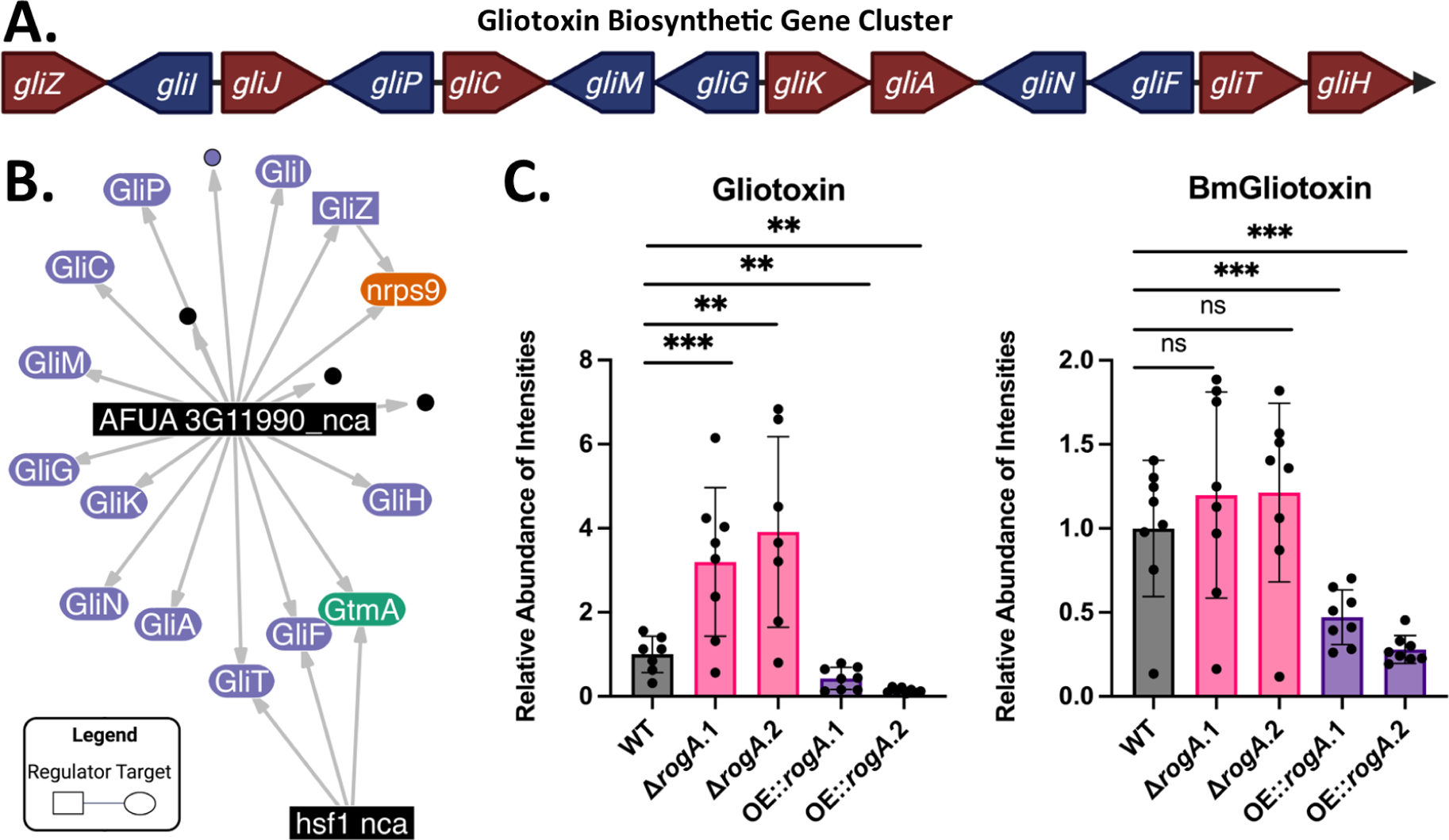
Validation of a novel predicted regulator from MERLIN-P-TFA and GRAsp in Gliotoxin regulation. A) Canonical gliotoxin biosynthetic gene cluster; colored by direction. B) MERLIN-P-TFA module 5349, visualized by GRAsp; nodes are colored by gene family with black nodes representing genes without common name in fungiDB. C) Production of Gliotoxin and bis(methylthio)gliotoxin in two independent *rogA* transformants of Δ*rogA* and OE::*rogA*. Relative abundance of gliotoxin and bis(methylthio)gliotoxin (BMgliotoxin) was measured by intensities of UHPLC-HRMS spectra. Results are the average relative abundance of eight repetitions ± standard deviation. Statistical analysis was performed using an unpaired two-sided t-test when comparing quantities to the relative abundance of the WT strain (* p<0.1, ** p<0.05, *** p<0.01). Samples were normalized with respect to the wild type of each author. Figure prepared with Biorender.com.

### MERLIN-P-TFA and GRAsp identify the AtfA transcription factor as a key component of the LCO signaling pathway in *A. fumigatus*

Fungi use a diverse array of signals to interact with their environment (82). One such prompt are lipo-chitooligosaccharides (LCOs), small signaling molecules consisting of a chitin backbone, an attached long-chain fatty acyl group, and various functional groups, such as sulfated and fucose groups (83–85). For decades, the scientific community thought LCOs were solely produced by plant symbiotic microbes, including nitrogen-fixing rhizobia and mycorrhizal fungi, as LCOs are critical for recognition by host plants. However, we have recently shown that LCOs are synthesized throughout the fungal kingdom and that many fungi respond to LCOs in a dose-dependent manner (33). In particular, *A. fumigatus* treatment with a specific endogenous LCO (LCO-IV = C:16, S) led to a significant reduction in hyphal branching (hypobranching) and an increased germination rate of conidia (33, 86).

To identify LCO signaling pathways leading to these phenotypes, Rush et al. performed RNA-seq analysis of LCO-IV (C:16, S) treated *A. fumigatus* cultures. However, the large number of differentially expressed genes (DEGs) made it challenging to identify key regulators to target with perturbation experiments. We used the node diffusion method in GRAsp to incorporate data from the differential expression analysis and prioritize regulators involved in LCO signaling. For each node, we associated it with an initial score based on the absolute value of the log fold change between *A. fumigatus* 30 minutes after treatment with LCOs or with a negative control, then applied laplacian kernel diffusion (λ = 10) (33). After diffusion we selected the top 10 regulators with at least 5 targets based on diffusion score (**Figure 6A**). The most connected of these regulators was the gene *atfA*. AtfA is a well-known bZIP transcription factor essential for stress tolerance of conidia and reactive oxygen intermediate resistance during invasive aspergillosis (57, 87). To test if *atfA* plays a role in fungal response to LCOs, we examined branching and germination rates of *A. fumigatus* treated or not with LCO-IV (C:16, S). **Figure 6B** shows that both △*atfA* and OE::*atfA* strains were unresponsive to LCO-IV compared to the wild-type. Furthermore, regardless of LCO treatment, the deletion of *atfA* led to increased hyphal branching (hyperbranching, top) and slow germination phenotypes (bottom). Reciprocally, the overexpression of *atfA* led to an extreme hypobranching phenotype even lower than the wild-type strain treated with LCO-IV. These results placed *atfA* as an essential component of the LCO signaling pathway and revealed its function in hyphal branching in *A. fumigatus*. *atfA* loss has been implicated in a germination reduction in *A. oryzae*, matching our observations with *A. fumigatus* (*88*).

**Figure 6.**
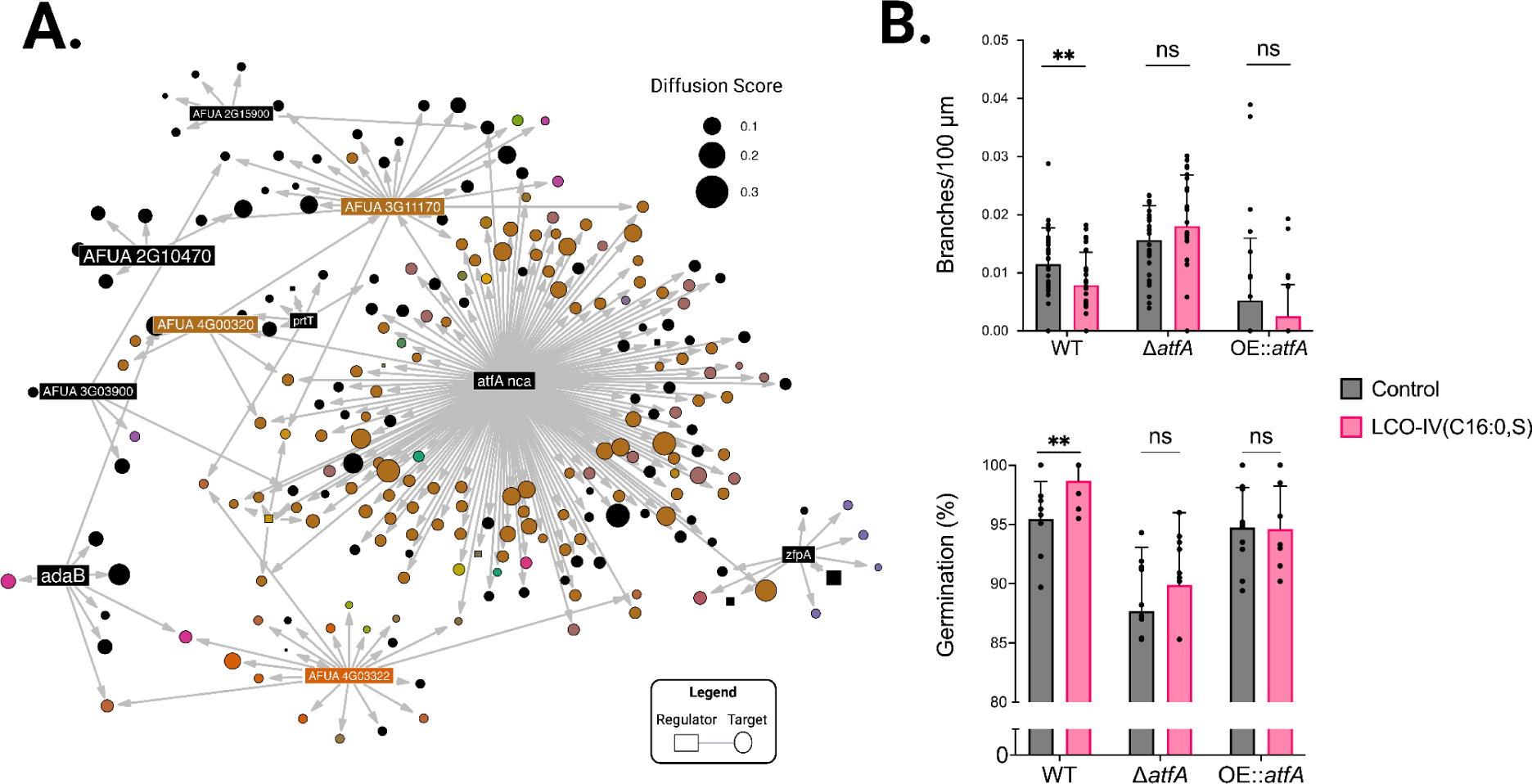
*atfA* is necessary for branching and germination responses to LCO-IV. A) Differentially expressed genes integrated with MERLIN-P-TFA network using network diffusion within GRAsp. Labeled regulators (rectangles) represent the top 10 regulators with at least 5 targets after diffusion of the absolute log fold change signal. Node size represents the score after diffusion. *AtfA* is shown as a key regulator with many differentially expressed targets (large node size). Genes are colored by module identity, and black nodes represent genes not assigned to a module. B) Secondary branching *of atfA* mutant hyphae 12 hours after treatment with LCO-IV (C:16, S) or negative control, 0.005% EtOH. C) Percent germination *of atfA* mutant conidia 10 hours after treatment with LCO-IV (C:16, S) or negative control, 0.005% EtOH. Statistical analysis was performed using an unpaired two-sided t-test (* p<0.1, ** p<0.05, *** p<0.01, NS, not significant). Germination and branching experiments were performed in triplicates. Figure prepared with Biorender.com.

### Steiner tree predicts novel components of the chitin synthesis pathway

The final feature of GRAsp we demonstrate in this work is application of the Steiner tree approximation algorithm which attempts to find connections between any set of genes regardless of whether they belong to the same module. As LCO biosynthesis is dependent on chitin availability and chitin is a critical component of the cell wall (89, 90), we set out to identify regulators targeting chitin synthase genes. In this example, we compare the networks that are generated by (i) modules that incorporate two genes encoding family one chitin synthases, *chsA*, and *chsG* (*91*) (**Figure 7A**) and (ii) the Steiner tree generated to connect these two nodes (**Figure 7B**). Figure prepared with Biorender.com.

**Figure 7.**
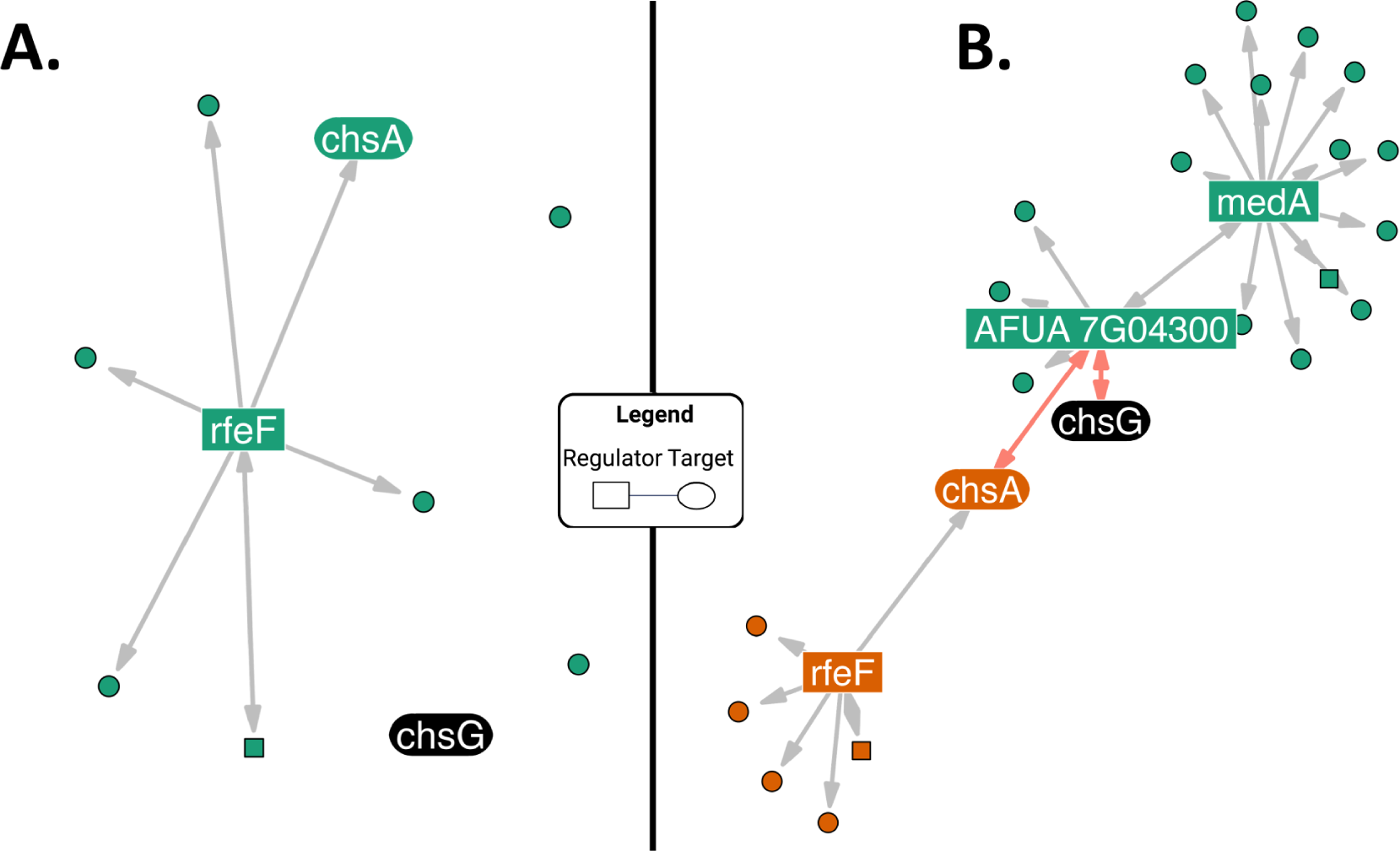
Predicted regulatory relationship of *chsA* and *chsG*. A) Network representation after searching for modules associated with *chsA* and *chsG*. B) An estimated Steiner tree generated to connect *chsA* and *chsB*. Red edges correspond to the Steiner tree while the remaining edges represent the genes in the associated modules. Ovals represent gene targets, while rectangles represent regulators. Genes are colored by module identity, and black nodes represent genes not assigned to a module. The Steiner tree approximation shows a previously uncharacterized relationship between *chsA* and *chsG*. Figure prepared with Biorender.com.

Notably, the Steiner tree network provides additional modules which capture biologically relevant information. No common regulator of *chsA* and *chsG* was identified when searching with the MERLIN module method. This can occur when genes are associated with independent modules without a common regulator. The Steiner tree method is useful for finding common regulators in these instances. After finding an approximate Steiner tree (red edges **Figure 7B**) between *chsA* and *chsG*, a common regulator AFUA_7G04300 was identified. AFUA_7G04300, an uncharacterized gene predicted to contain a RhoGAP and Fes/CIP4 domain, regulates not only *chsA* and *chsG* but is also predicted to regulate *gfa1*, a putative glutamine-fructose-6-phosphate transaminase. This was particularly interesting because this enzyme would theoretically catalyze the first reaction in the chitin synthesis process: the formation of glucosamine 6-phosphate (89). This predicted regulatory relationship would not have been identified without the Steiner search algorithm.

AFUA_7G04300 was also implicated in broader cell wall activities, as it was predicted to regulate AFUA_5G13090, a putative alpha-1,2-mannosyltransferase and cell wall biosynthesis gene (92). AFUA_7G04300 is also connected to *rgd1*, a putative Rho-GTPase activating protein whose homolog in *Saccharomyces cerevisiae* has been implicated in cell wall integrity signaling(93). The modules containing the Steiner tree also connected *rfeF* to *chsA.* RfeF is a transcription factor regulated by CrzA, itself a transcription factor known to bind to promoters of both *chsA* and *chsG* and thus is tied into chitin synthesis (94). The *chsC/chsG* Steiner tree presents a plausible connection between chitin regulatory pathways that can be further validated and explored.

## DISCUSSION

In October 2022, the World Health Organization (WHO) published its first-ever report of priority fungal pathogens (95). *A. fumigatus* - along with *Cryptococcus neoformans, Candida albicans*, and *Candida auris* - was placed in the critical priority group(95). This placement was due to a substantial increase in severe aspergillosis cases, burgeoning mortality rates, co-infections with COVID and other respiratory pathogens, and increasing antifungal resistance of this mold(95). Accordingly, and accompanying this recognition of *A. fumigatus* as a critical infectious agent, laboratories worldwide have focused on elucidating the genes and molecules that make *A. fumigatus* a successful pathogen (96, 97). Although a large number of RNA-seq experiments have been performed to understand the mode of virulence of *A. fumigatus* and, to ultimately identify new targets for antifungal development (33, 34, 98, 99), these efforts alone have not fully revealed the regulatory networks and pathways required for the pathogenicity of this fungus. For example, although it has long been known that gliotoxin is a virulence determinant of invasive aspergillosis, a full understanding of the signaling networks promoting the synthesis of this toxin remains obscure. Our goal in this study was to leverage the wealth of existing RNA-seq datasets available for this fungus to construct a genome-scale GRN and make it easily accessible to the research community at large and, further, to integrate existing knowledge to predict regulators important in regulating fungal development and virulence factors.

By leveraging publicly available gene-expression data of multiple strains of *A. fumigatus*, MERLIN-P-TFA was able to reconstruct, to our knowledge, the first comprehensive gene regulatory network of the organism. We developed GRAsp, a user-friendly web portal, to query this inferred GRN with individual genes and gene sets to predict novel components of diverse biological pathways using one of three integrated features: Regulatory Modules, Steiner trees, or node diffusion. Here we used regulatory modules to successfully recapitulate previously known pathways such as SrbA regulation of genes involved in ergosterol biosynthesis, nitrate assimilation, and heme biosynthesis during hypoxia; HapX regulation of genes involved in the synthesis and transport of siderophores; ZafA regulation of three zinc transporters *zrfA*, *zrfB*, and *zrfC* and FapR regulation of genes involved in the synthesis of the secondary metabolites, pseurotin and fumagillin.

The successful recovery of several known metabolic pathways using MERLIN-P-TFA and GRAsp led us to test the ability of GRAsp to identify unknown components regulating the modules. For our example, we choose to investigate gliotoxin regulation. Gliotoxin was isolated from several fungi in the 1940s, where it was quickly found to exhibit strong antifungal activity(100). Consideration of this metabolite for such use declined with the finding of its general eukaryotic toxicity. Identification of the *A. fumigatus* gliotoxin gene cluster allowed for molecular characterization and *gli* gene deletions, all leading to the conclusion that gliotoxin is an exacerbating factor in invasive aspergillosis (101, 102). Further, these studies showed how *A. fumigatus* protected itself from its toxin, dependent on several enzymes, including GliT and GtmA, that modified/prevented the toxic disulfide bridge in the gliotoxin molecule. Although the *gli* BGC regulator GliZ was characterized in 2006, it is unknown how *gliZ* is regulated (80). The input of *gliZ* into the GRAsp tool revealed a module predicted to be regulated by a gene coding for a putative transcription factor AFUA_3G11990, now named *rogA* (**Figure 5A**). This module contained all of the *gli* BGC genes as well as the trans-located *gtmA* gene involved in self-protection. Examination of *rogA* (AFUA_3G11990) mutants revealed that this putative transcription factor is a negative regulator of gliotoxin production (**Figure 5B**).

The accuracy of MERLIN-P-TFA and GRAsp predicted regulators was further supported by the identification of AtfA as a key transcription factor required for the developmental response of *A. fumigatus* to LCOs. These LCO signals are one of two endogenous signals we identified to regulate lateral branching in *A. fumigatus* (*33, 34*). Filamentous fungi such as *A. fumigatus* must balance hyphal extension with lateral branching for optimal colony and invasive growth. Understanding the GRNs transmitting these signals has the potential for future therapies to control the growth of *A. fumigatus.* We used node diffusion to discover that AtfA is required for *A. fumigatus* to respond to LCOs, thus uncovering a previously unknown mechanism of LCO signaling pathway(s) in fungi.

Finally, we use Steiner trees to predict AFUA_7G04300 as a new regulator of two chitin synthases, *chsA* and *chsG*. While it needs to be validated experimentally, the prediction is further backed up by the other predicted targets of AFUA_7G04300, which are involved in cell wall integrity and possibly the synthesis of chitin precursors.

Our work shows the strength of our gene regulatory network as a powerful resource for generating new hypotheses. Still, at 80% confidence, the network only captures ∼30% of all genes in the *A. fumigatus* genome, with 2,940 genes out of the total of 9,859 genes of this fungus into high confidence modules (**Figure 2**). One consequence is that some well-described pathways, like the BrlA-mediated conidiation pathway (103), are not represented in GRAsp. Furthermore, MERLIN-P-TFA and GRAsp did not always pick up every single gene in the recovered pathways, which we postulate is due to the limitation of the number of RNA-seq datasets used in this study.

However, as new expression datasets become publicly available, we can incorporate them into our framework to refine existing pathways and potentially uncover new pathways. Beyond RNA-seq, other complementary omic data (e.g., ChiP-seq or ATAC-seq) can also be incorporated into the MERLIN-P-TFA framework, increasing our network’s coverage and accuracy. Follow up transcriptomic and epigenomic studies guided by the prioritized regulators using GRAsp could more efficiently improve the quality and coverage of our GRNs. To facilitate community access, GRAsp is available as a free online visualization tool that can be used to examine this GRN by individual researchers and generate model-driven testable hypotheses. Such model-driven experiments would additionally provide a quantitative assessment of the network’s quality and enable us to determine the most useful experiments towards constructing a comprehensive GRN for *A. fumigatus*. We believe that our inferred GRN and associated GRAsp tool will be a useful resource for identifying the key genes and pathways that underlie the pathogenic traits of *A. fumigatus*.

## AVAILABILITY

Data preprocessing was performed utilizing Trimmomatic (v0.32), FastQC (v0.11.9), MultiQC, RSEM (v1.2.11), Bowtie2 (v2.2.0), and MATLAB (r2017b). Data preprocessing scripts and processed data can be found https://github.com/Roy-lab/merlin-preprocess.

Network inference was performed utilizing the MERLIN-P-TFA package (https://github.com/Roy-lab/MERLIN-P-TFA, Siahpirani et al 2023 in preparation). MERLIN-P-TFA is dependent on the network component analysis package, EstimateNCA (https://github.com/Roy-lab/EstimateNCA), and the network inference algorithm, MERLIN-P (https://github.com/Roy-lab/merlin-p). After generating the network, analysis of the network structure was performed using the MERLIN-Auxiliary package (https://github.com/Roy-lab/merlin-auxillary).

The GRAsp RShiny app was constructed using the MERLIN-VIZ package (https://github.com/Roy-lab/MERLIN-VIZ). The GRAsp branch contains net_data.Rdata which stores all network data. A wrapper function used to make all networks figures in this paper is also provided. The GRAsp RShiny app is hosted at https://grasp.wid.wisc.edu/ and is a publicly available resource for hypothesis generation. GRAsp was constructed using the tidyverse package suite (v2.0.0), tidygraph (v1.2.3), pracma (v2.4.2) and igraph (v1.4.1). Network visualization in GRAsp utilizes networkD3 (v0.4), ggraph (v2.1.0), and RColorBrewer (v1.1-3). Tables in GRAsp utilize the DT package (v0.27).

## ACCESSION NUMBERS

The RNA-seq dataset generated in-house for this study has been deposited with GEO under accession number GSE231238. It can be accessed with reviewer’s token wxoziaowndsbdwr. The remaining accession number for the publically available RNA-seq datasets utilized in this study are contained in supplementary **Table S1**.

## SUPPLEMENTARY DATA

The following Supplementary Files are available at NAR online. **GRAsp_supplementary_text.pdf** provides the Supplementary Text. **S1_regulators.xlsx** presents the list of regulators.

**S2_prior_network.xlsx** presents the prior network.

**S3_module_details.xlsx** presents MERLIN consensus module assignment and GO term enrichment per module.

## Supporting information

S1_regulators.xlsx

S2_prior_network.xlsx

S3_module_details.xlsx

