## Supplementary Text for "A network-based model of *Aspergillus fumigatus* elucidates regulators of development and defensive natural products of an opportunistic pathogen"

### Table of Contents

**Supplementary Tables**


- Supplementary Table S1. RNAseq datasets used to create GRAsp
- Supplementary Table S2. Fungal strains used in this study
- Supplementary Table S3. PCR primers for this study


**Supplementary Figures**


- Supplementary Figure S1. The number of regulatory genes curated from five different sources
- Supplementary Figure S2. The rationale behind the construction of the prior network based on sequence motifs
- Supplementary Figure S3. Confirmation of rogA mutants
- Supplementary Figure S4. A heatmap of the zero mean quantile-normalized data of genes in module 5349
- Supplementary Figure S5. Principal components of samples SI1-SI8.
- Supplementary Figure S6. Principal components of samples SI9-SI13.
- Supplementary Figure S7. Principal components of samples SI14-SI18.
- Supplementary Figure S8. Reference SI 2 (PRJEB2987) sample correlation.
- Supplementary Figure S9. Reference SI 2 (PRJEB2987) expression heatmaps.


**Supplementary Methods**


- Supplementary Method S1. Descriptions of the previously characterized TF binding site sequence motifs

### Supplementary Tables

**Supplementary Table S1. RNAseq datasets used to create GRAsp.** Dataset ID refers to Bioproject Accession. Sample No. refers to the number of samples from the dataset used to create GRAsp. S.I No., Serial Number.

| Sl. No. | Dataset ID | Year | WIldtype Strain | Description | Sample No. | Reference |
|---|---|---|---|---|---|---|
| 1 | PRJEB3185 | 2012 | CEA17 | WT and Δ*mpkA* | 6 | (Müller et al. 2012) |
| 2 | PRJEB2987 | 2014 | Af293 | WT and Δ*gliT* | 15 | (O'Keeffe et al. 2014) |
| 3 | PRJNA390719 | 2017 | Af293 | WT, Δ*rax1*, Δ*rgsD* | 6 | (Choi et al. 2019, Kim et al. 2019) |
| 4 | PRJNA508764 | 2018 | Af293 | WT, Δ*atrR, atrR*-3X HA ,hspA-*atrR* | 8 | (Paul et al. 2019) |
| 5 | PRJNA554811 | 2019 | ATCC 46645 | WT and cytomegalovirus coinfection of human dendritic cells | 16 | (Seelbinder et al. 2020) |
| 6 | PRJNA558954 | 2019 | ATCC 46645 | WT exposure to human dendritic cells (2,4,6 h). MOI of 1,0,5 | 9 | (Seelbinder et al. 2020) |
| 7 | PRJNA144647 | 2011 | CEA10 | Treatment of hypoxia or normoxia (12,24,36 h) | 4 | (Losada et al. 2014) |
| 8 | PRJNA240324 | 2014 | CEA17 | WT and Δ*akuB* exposure to humidimycin | 12 | (Valiante et al. 2015) |
| 9 | PRJNA240892 | 2014 | A1163 | Caspofungin exposure of WT and Δ*akuB* (0,0.5,1,4,8 h); Δ*mpkA* and Δ*sakA* (0,1,4 h) | 53 | (Altwasser et al. 2015) |
| 10 | PRJNA241401 | 2014 | CEA10 | WT exposure to hypoxic or normoxic conditions (0,15,30 min) | 12 | (Hillmann et al. 2014) |
| 11 | PRJNA376829 | 2017 | Af293 | WT grown on sugarcane bagasse or fructose | 6 | (de Gouvêa et al. 2018) |

| | | | | | | |
|---|---|---|---|---|---|---|
| 12 | PRJNA399754 | 2017 | CEA10/Af293 | WT infection of A549 type II pneumocyte cell line (6,16 h) | 24 | (Watkins et al. 2018) |
| 13 | PRJNA408076 | 2017 | AFIR964/AFIR974 | WT germination (0,2,4,6,8 h) | 20 | (Baltussen et al. 2018) |
| 14 | PRJNA482512 | 2018 | Af155-40<br>Af130–14<br>Af147–03 | WT exposure to itraconazole (30,60,120,240 min) | 30 | (Hokken et al. 2019) |
| 15 | PRJNA622251 | 2020 | CEA17 | WT and Δ*fhdA* exposure to caspofungin (48 h) | 24 | (Valero et al. 2020) |
| 16 | PRJNA642658 | 2020 | Af293 | WT exposure to LCO-IV (30 min and 2 h) | 16 | (Rush et al. 2020) |
| 17 | PRJNA658306 | 2020 | Af293 | WT and Δ*ppoA* exposure to 5,8-diHODE (30 min and 2 h) | 16 | (Niu et al. 2020) |
| 18 | GSE231238 | 2023 | Af293 | WT, OE::*zfpA*, and Δ*zfpA* at 0 hours and after 16 hours of shaking incubation | 24 | This study |

**Supplementary Table S2. Fungal strains used in this study**

| Name | Genotype | Reference |
|---|---|---|
| CEA17 KU80 | pyrG1, ∆akuB::pyrG, pyrG1 | (da Silva Ferreira et al. 2006) |
| TAJS1.7 | pyrG1, ∆akuB::pyrG, pyrG1, ∆AFUB_037850::pyrG | (Wiemann et al. 2017) |
| TAJS2.2 | pyrG1, ∆akuB::pyrG, pyrG1, ∆AFUB_064280::pyrG, gpdA(p)::AFUB_037850::argB | (Wiemann et al. 2017) |
| TFYL80.1 | *A.fumigatus fumiargB; ΔnkuA::mluc; pyrG1; argB1* | (Lim et al. 2018) |
| TSCP2.1 | *A.fumigatus fumiargB; ΔnkuA::mluc; pyrG1; argB1; ΔrogA::para_pyrG* | This study |
| THWS25.1 | *A.fumigatus fumiargB; ΔnkuA::mluc;pyrG1;argB1;argB1;para_pyrG::gpdA(P)::rogA* | This study |
| TFYL81.5 | *A. fumigatus pyrG1; argB1; ∆akuA::mluc; fumiargB; fumipyrG* | (Throckmorton et al. 2016) |
| TJW213.1 | *A. fumigatus pyrG1; argB1; ∆akuA::mluc; fumiargB; ∆zfpA::parapyrG* | (Schoen et al. 2023) |
| TJW214.2 | *A. fumigatus pyrG1; argB1; ∆akuA::mluc; fumiargB; parapyrG::gpdA(p)::zfpA* | (Schoen et al. 2023) |

**Supplementary Table S3. PCR primers for this study**

| Name | Sequence (5'-3') | Use |
|---|---|---|
| A. para pyrG For. | GTCGACGGTATCGATAAGCTTG | *ΔrogA* |
| A. para pyrG Rev | ATTCGACAATCGGAGAGGCTGC | *ΔrogA* |
| del_AFUA3G11990_3'F | CTGTCGCTGCAGCCTCTCCGATTGTCGAAT GTACGTTCCCAGTACAACTACTGAGCC | *ΔrogA* |
| del_AFUA3G11990_3'R | CATCCACTGAATCCAGTCGTCG | *ΔrogA* |
| del_AFUA3G11990_5'F | AGCAGCCGTTTGAGAGTCATGC | *ΔrogA* |
| del_AFUA3G11990_5'R | CGATATCAAGCTTATCGATACCGTCGACCGTT GGCGCAAAGGCTCATGCAAGG | *ΔrogA* |
| OE_AFUA_3G11990_5F | CGAGGCTTAGGCTTCTTCGAAC | OE::*rogA* |
| OE_AFUA3G11990_5R | CCTCTCGGGCCATCTGTTCGTATAAGCTTCT CGACGATGACGTTGGCGCAAAGGC | OE::*rogA* |
| OE_AFUA_3G11990_3F | CTACCCCGCTTGAGCAGACATCACCATATGG CCGCCAATTCCAGTCCGTTC | OE::*rogA* |
| OE_AFUA_3G11990_3R | CCGGAGGGGTATCATAGATTCG | OE::*rogA* |
| PyrG+gpdA(P) F | CGTAATACGACTCACTATAGGGC | OE::*rogA* |
| PyrG+gpdA(P) R | GGTGATGTCTGCTCAAGCGGG | OE::*rogA* |

### Supplementary Figures

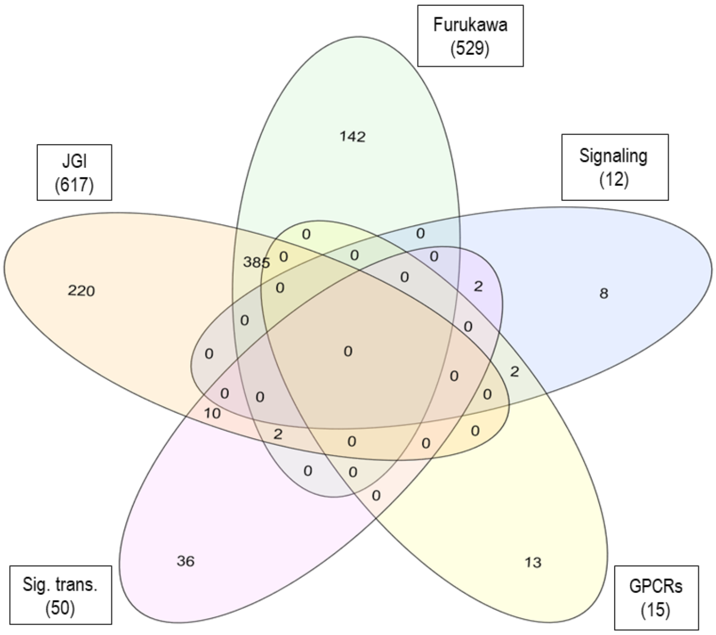


| Source | No. of genes |
|---|---|
| JGI MyCoCosm | 617 |
| Furukawa et al. | 529 |
| Signaling genes | 12 |
| G-protein coupled receptors (GPCRs) | 15 |
| Signal transduction genes | 50 |
| *All combined* | *820* |

**Supplementary Figure S1. The number of regulator genes curated from five different sources.** The Venn diagram presents the number of unique regulators contributed by each source. For example, we curated 617 regulators from the JGI MyCoCosm database (Grigoriev et al. 2014). However, only 220 of them are unique to this database i.e., they were not present in the sublists of regulators collected from other sources. Combining all the sources resulted in a list of 820 distinct regulators. Full list of regulators can be found in **Supplementary File S1_regulators.xlsx**.

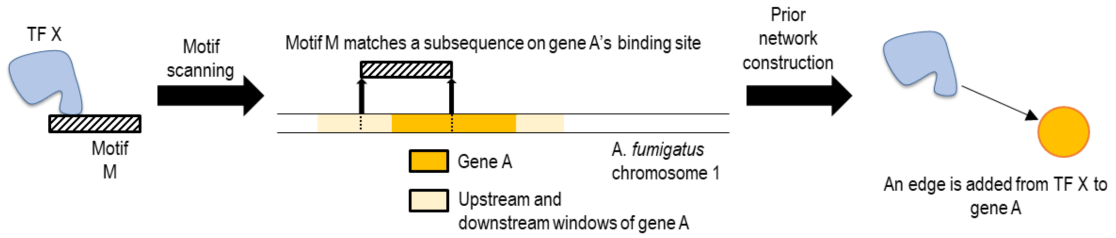


**Supplementary Figure S2. The rationale behind the construction of the prior network based on sequence motifs.** If transcription factor (TF) X can bind to DNA-sequence motif M, we scanned the A. *fumigatus* reference genome to find matches for motif M using the pwmmatch.exact.r of PIQ package from Sherwood et al. 2014. We defined the binding region of a gene to be 10 Kbp upstream of the gene through 1 Kbp downstream of the gene. If a match was found on the binding region (such as the promoter) of Gene A in chromosome 1, we added an edge from TF X to Gene A in the prior network.

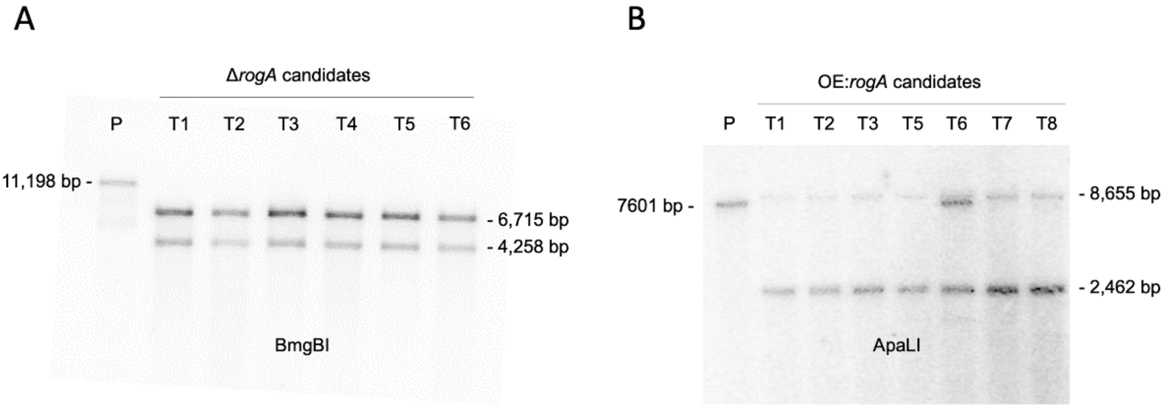


**Supplementary Figure S3. Confirmation of *rogA* mutants.** A) Southern confirmation of Δ*rogA* mutants. Genomic DNA was digested by BmgBI (Parent: P, 11198 bp; Transformants: T, 4258 and 6715 bp). B) Southern blot confirmation of OE::*rogA* mutants. Genomic DNA was digested by ApaLI (Parent: P, 7601 bp; Transformants: T, 2462 and 8655 bp).

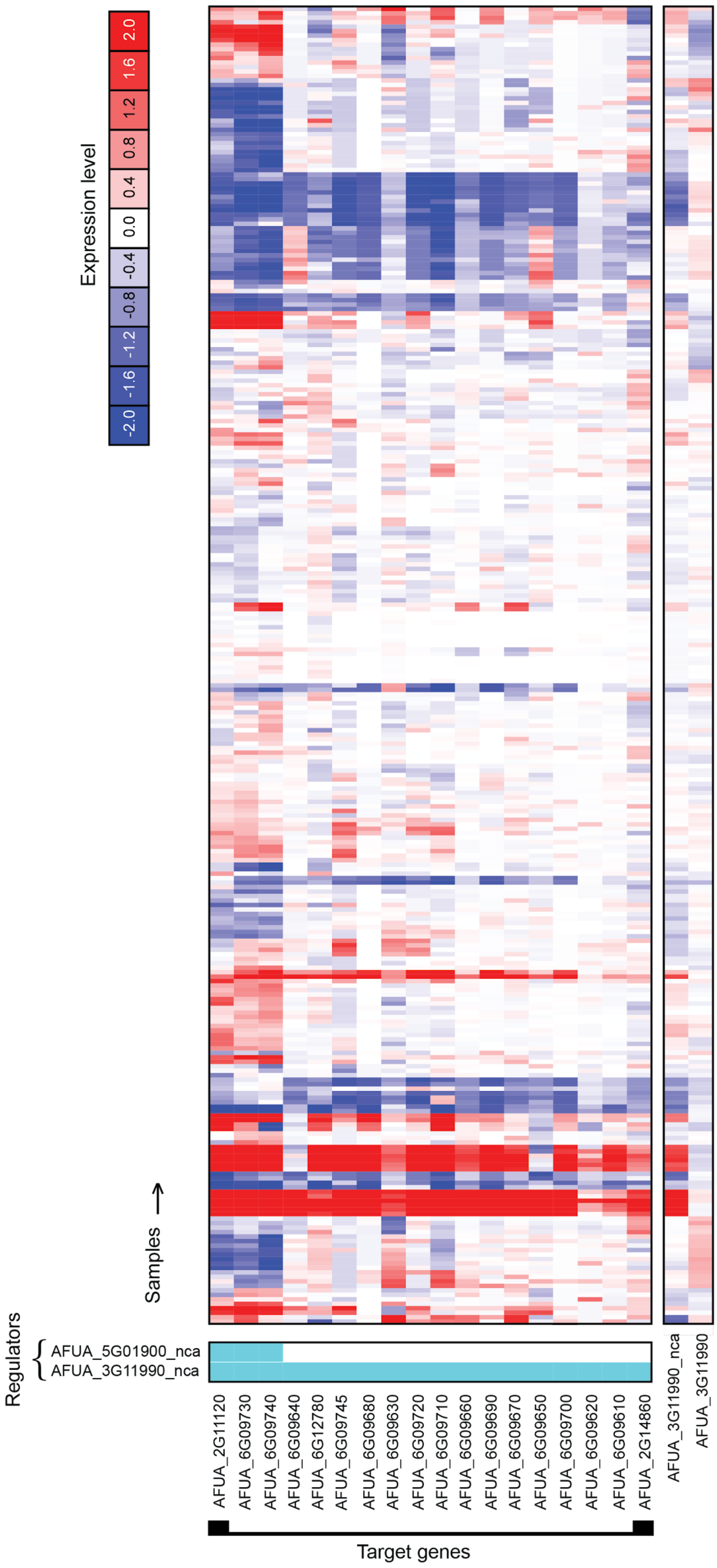


**Supplementary Figure S4. A heatmap of the zero mean quantile-normalized data of genes in module 5349.** Systematic names for genes are on rows and samples are on columns. The profiles of regulators are depicted below genes in the modules, the last two rows. Regulator-target interactions are also labeled in the first two columns with light blue squares. AFUA_3G11990 is anti-correlated with many of the genes in the module, suggesting that it is a repressor of gliotoxin biosynthesis. The transcription factor activity of AFUA_3G11990, AFUA_3G11990_nca, is also depicted. The TFA profile is correlated with the gliotoxin genes within the module.

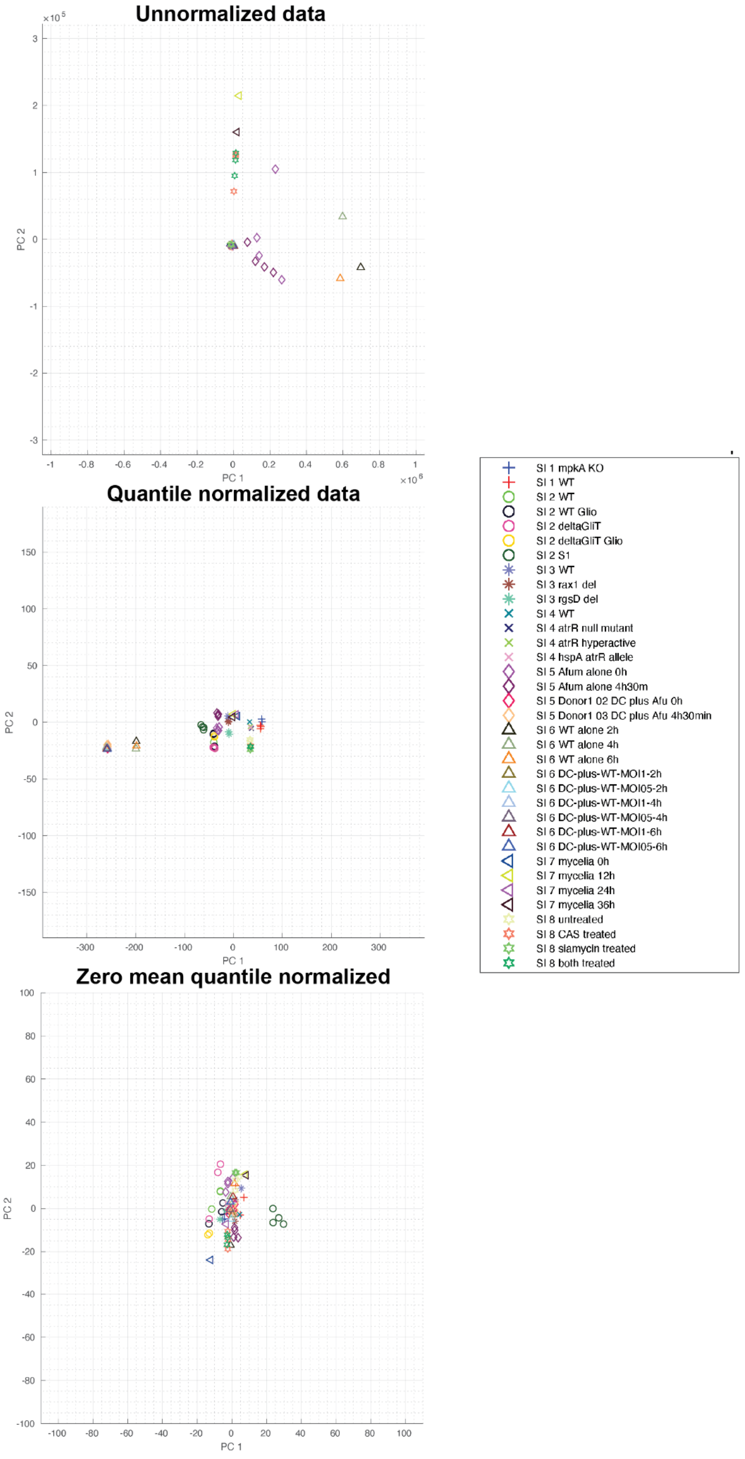


**Supplementary figure S5. Principal components of samples SI 1-SI 8.** Principal component scores of all samples were calculated simultaneously. Displayed are the PCA scores of samples from SI 1-SI 8. PCA of unnormalized and quantile normalized data demonstrates batch effects, that is, samples are clustered together by batch (shape) regardless of the experimental condition (color). After zero mean normalization, the batch effect is largely removed. Samples are clustered by similar experimental conditions, regardless of batch.

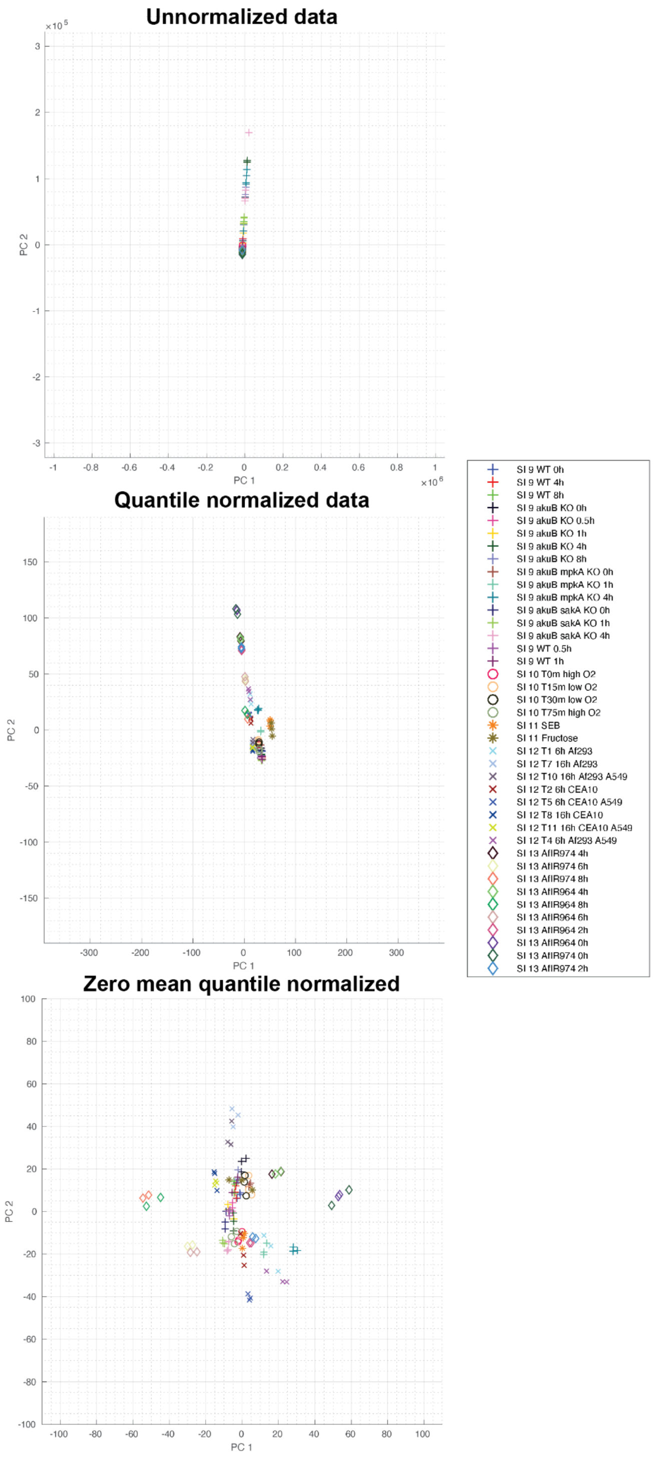


**Supplementary figure S6. Principal components of samples SI 9-SI 13.** Principal component scores of all samples were calculated simultaneously. Displayed are the PCA scores of samples from SI 9-SI 13. PCA of unnormalized and quantile normalized data demonstrates batch effects. Samples are clustered together by batch (shape) regardless of the experimental condition (color). After zero mean normalization, the batch effect is largely removed. Samples are clustered by similar experimental conditions, regardless of batch.

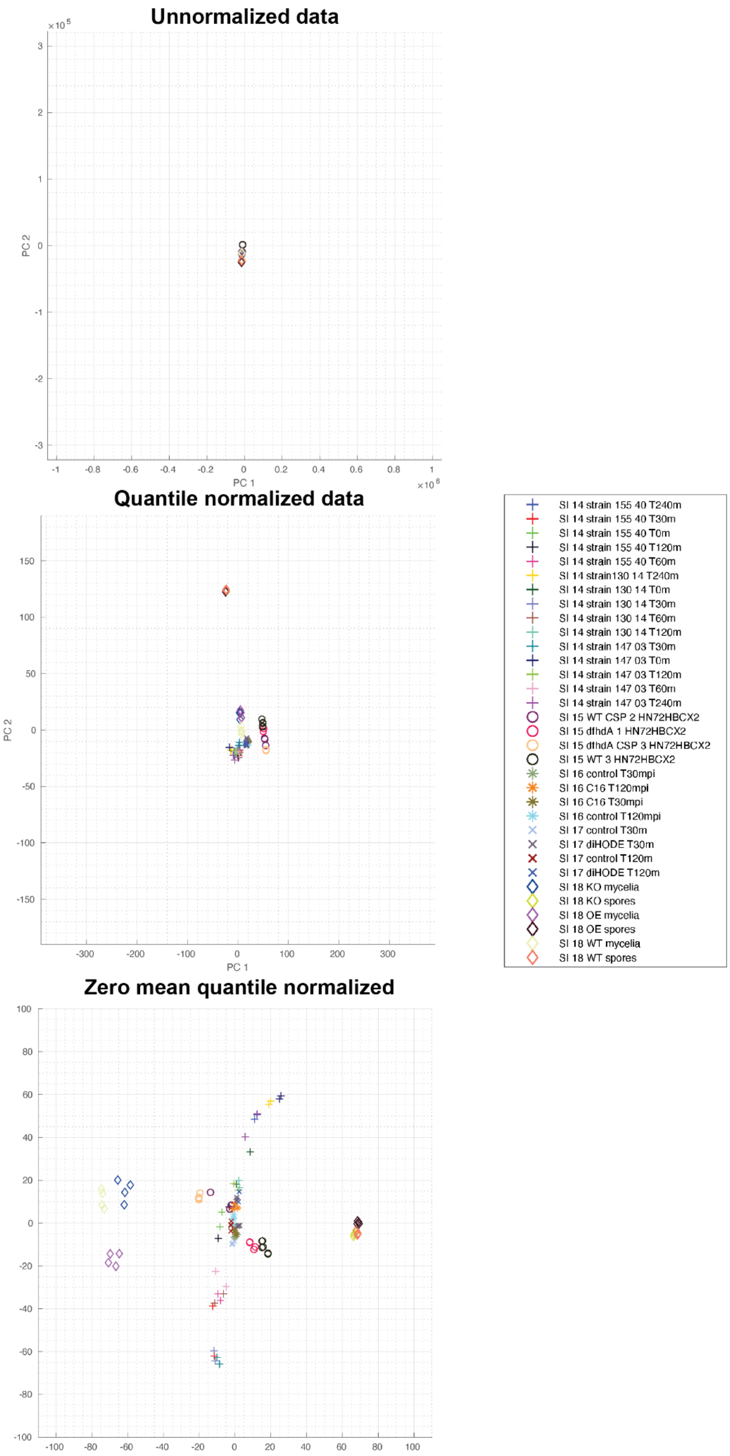


**Supplementary figure S7. Principal components of samples SI 14-SI 18.**Principal component scores of all samples were calculated simultaneously. Displayed are the PCA scores of samples from SI 14-SI 18. PCA of unnormalized and quantile normalized data demonstrates batch effects. Samples are clustered together by batch (shape) regardless of the experimental condition (color). After zero mean normalization, the batch effect is largely removed. Samples are clustered by similar experimental conditions, regardless of batch.

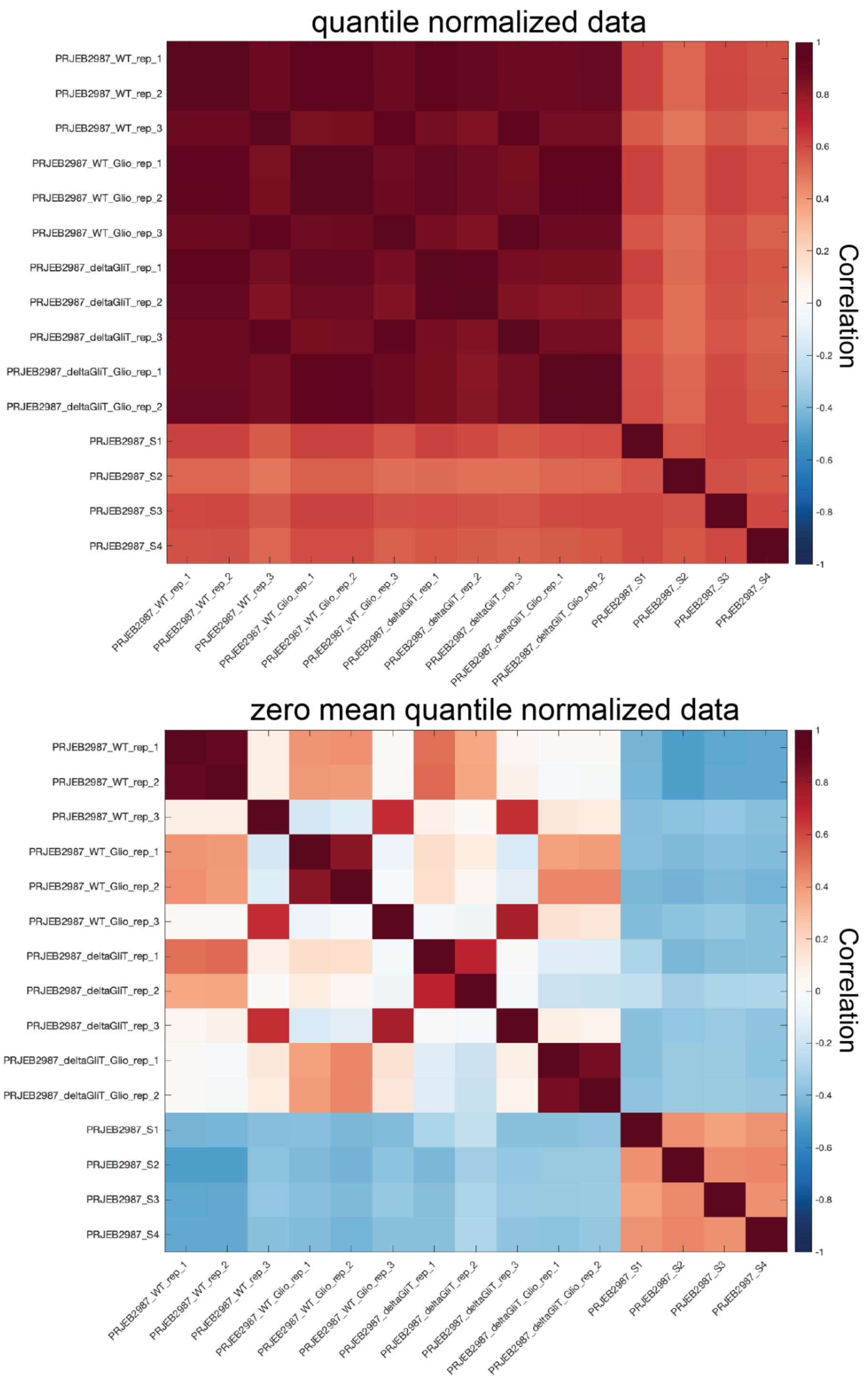


**Supplementary Figure S8: Reference SI 2 (PRJEB2987) sample correlation:** Samples of SI 2 are all strongly correlated prior to batch correction by zero-mean normalization. After zero-mean correction replicates 1 and 2 of each condition are strongly correlated, e.g. WT rep1 and 2. Rep 3 samples are strongly correlated regardless of condition and were removed prior to network inference.

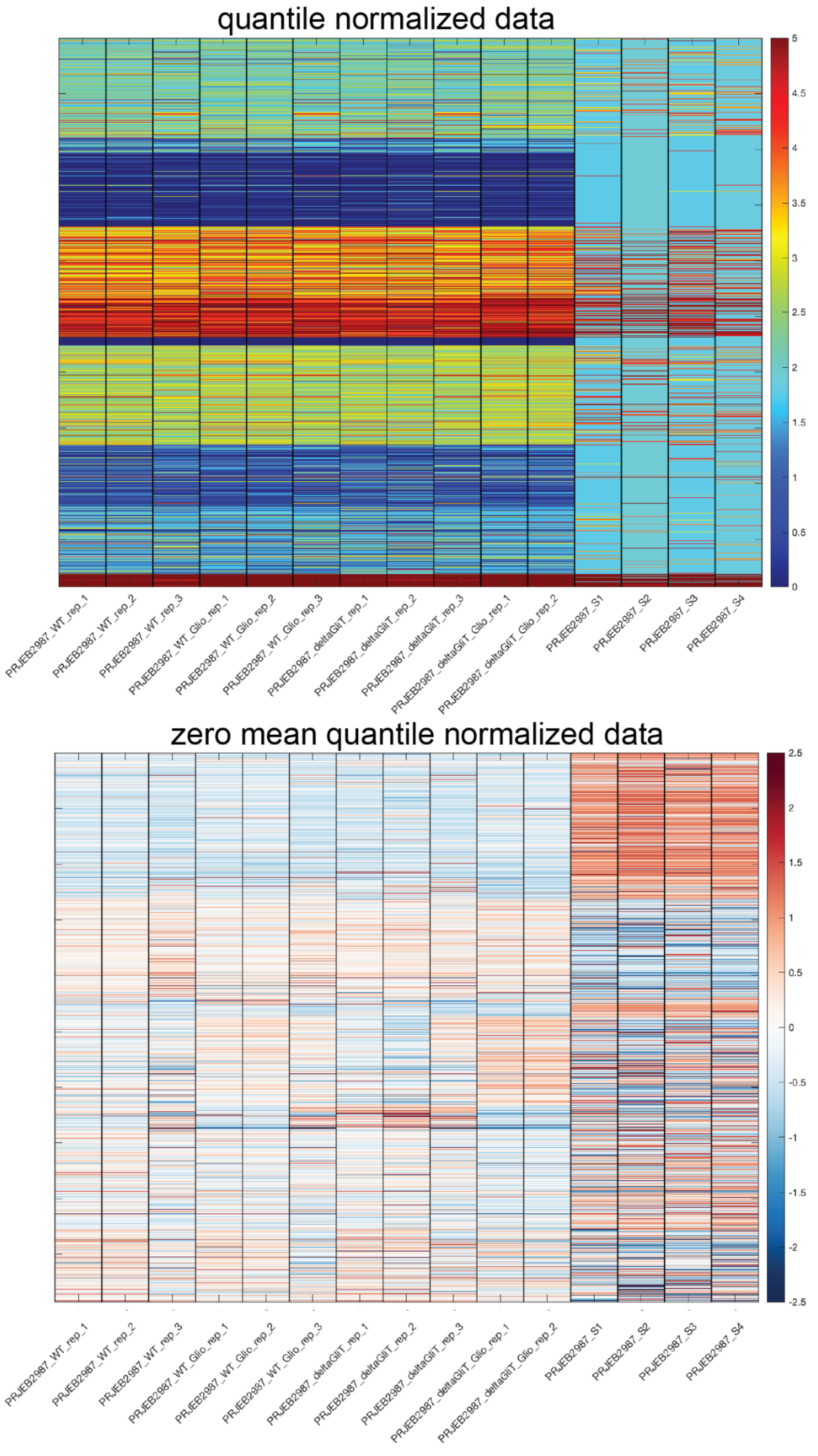


**Supplementary Figure S9: Reference SI 2 (PRJEB2987) expression heatmaps:** Expression of genes per sample before and after zero-mean normalization. Expression of SI 2 genes in quantile normalized data are increased relative to other samples (~1.75). These samples were removed prior to network inference because of this inconsistency.

### Supplementary Methods

#### Supplementary Method S1. Descriptions of the previously characterized TF binding site sequence motifs

In this section, we describe the previously characterized TF binding site sequence motifs used in this study. There are a total of four such motifs corresponding to two TFs – somA and xanC – as described below.

**TF name: somA**

Protein: Transcriptional activator somA
(ORF) GENEID: AFUA_7G02260 (source: https://www.uniprot.org/uniprot/Q4WAR8 )
The following motif preferences were recently characterized by Chen et al. [DOI: https://doi.org/10.1128/mBio.02329-20].
Motif 1: GTACTCCGTAC
Motif 2: RTGGBMTGATS
Motif 3: CCTMCAGAGCAG
Note 1: IUPAC symbols for bases are used
(Reference for the IUPAC symbols: https://www.bioinformatics.org/sms/iupac.html ).
R = A/G
B = C/G/T
M = A/C
S = C/G
Note: We assumed that the probability of having a base other than the base(s) given for a particular base position is low.

**TF name: xanC**

Protein: Xanthocillin biosynthesis cluster transcription factor xanC
(ORF) GENEID: AFUA_5G02655 (source: https://www.uniprot.org/uniprot/A4DA05)
The following motif preference was recently characterized by Wang et al. [DOI: https://doi.org/10.1128/mBio.01399-21]
The motif is 5'-(A)GTCAGC(A)-3' (motif 4)
The parenthesized "(A)" implies that we are not certain whether the binding motif contains A's or not. All of the xan promoters had 5'-AGTCAGCA-3' but there could be variations. Hence, we decided to exclude the parenthesized parts and use 5'-GTCAGC-3' as the motif.

XanC is the bZip transcription factor of the xanthocillin biosynthetic gene cluster. We conjectured that XanC might be a regulator of the copper homeostasis genes. That motivated us to include XanC's binding motifs in this study.

**Converting sequence motifs into position weight matrices (PWMs)**

For each of the four motifs, we produced a position weight matrix (PWM) by using 0.9 to indicate a high probability. Let us illustrate with an example. Suppose, a particular base position in a given motif has a high probability for G, and very low probabilities for the remaining letters. Then G will be assigned a probability value of 0.9 and 0.1 will be equally distributed between the remaining letters. Similarly, if more than one letter has a high probability for a particular position, then 0.9 will be equally distributed among them. Again, 0.1 will be equally distributed between the remaining letters.Thus, we produced the PWMs of the previously characterized motifs; the PWMs are presented below.

The position weight matrix of motif 1 (GTACTCCGTAC):

| Position | A | C | G | T |
|---|---|---|---|---|
| 1 | 0.1 | 0.1 | 0.9 | 0.1 |
| 2 | 0.1 | 0.1 | 0.1 | 0.9 |
| 3 | 0.9 | 0.1 | 0.1 | 0.1 |
| 4 | 0.1 | 0.9 | 0.1 | 0.1 |
| 5 | 0.1 | 0.1 | 0.1 | 0.9 |
| 6 | 0.1 | 0.9 | 0.1 | 0.1 |
| 7 | 0.1 | 0.9 | 0.1 | 0.1 |
| 8 | 0.1 | 0.1 | 0.9 | 0.1 |
| 9 | 0.1 | 0.1 | 0.1 | 0.9 |
| 10 | 0.9 | 0.1 | 0.1 | 0.1 |
| 11 | 0.1 | 0.9 | 0.1 | 0.1 |

The position weight matrix of motif 2 (RTGGBMTGATS):
R = A/G; B = C/G/T; M = A/C; S = C/G.

| Position | A | C | G | T |
|---|---|---|---|---|
| 1 | 0.45 | 0.05 | 0.45 | 0.05 |
| 2 | 0.1 | 0.1 | 0.1 | 0.9 |
| 3 | 0.1 | 0.1 | 0.9 | 0.1 |
| 4 | 0.1 | 0.1 | 0.9 | 0.1 |
| 5 | 0.1 | 0.3 | 0.3 | 0.3 |
| 6 | 0.45 | 0.45 | 0.05 | 0.05 |
| 7 | 0.1 | 0.1 | 0.1 | 0.9 |
| 8 | 0.1 | 0.1 | 0.9 | 0.1 |
| 9 | 0.9 | 0.1 | 0.1 | 0.1 |

| | | | | |
|---|---|---|---|---|
| **10** | 0.1 | 0.1 | 0.1 | 0.9 |
| **11** | 0.05 | 0.45 | 0.45 | 0.05 |

The position weight matrix of motif 3 (CCTMCAGAGCAG):
R = A/G; B = C/G/T; M = A/C; S = C/G.

| **Position** | **A** | **C** | **G** | **T** |
|---|---|---|---|---|
| **1** | 0.1 | 0.9 | 0.1 | 0.1 |
| **2** | 0.1 | 0.9 | 0.1 | 0.1 |
| **3** | 0.1 | 0.1 | 0.1 | 0.9 |
| **4** | 0.45 | 0.45 | 0.05 | 0.05 |
| **5** | 0.1 | 0.9 | 0.1 | 0.1 |
| **6** | 0.9 | 0.1 | 0.1 | 0.1 |
| **7** | 0.1 | 0.1 | 0.9 | 0.1 |
| **8** | 0.9 | 0.1 | 0.1 | 0.1 |
| **9** | 0.1 | 0.1 | 0.9 | 0.1 |
| **10** | 0.1 | 0.9 | 0.1 | 0.1 |
| **11** | 0.9 | 0.1 | 0.1 | 0.1 |
| **12** | 0.1 | 0.1 | 0.9 | 0.1 |

The position weight matrix of motif 4 (GTCAGC):

| **Position** | **A** | **C** | **G** | **T** |
|---|---|---|---|---|
| **1** | 0.1 | 0.1 | 0.9 | 0.1 |
| **2** | 0.1 | 0.1 | 0.1 | 0.9 |
| **3** | 0.1 | 0.9 | 0.1 | 0.1 |
| **4** | 0.9 | 0.1 | 0.1 | 0.1 |
| **5** | 0.1 | 0.1 | 0.9 | 0.1 |
| **6** | 0.1 | 0.9 | 0.1 | 0.1 |
